# Acute activation of Gq-signaling in islet macrophages inhibits β-cell insulin secretion through AMPK-sphingolipid axis

**DOI:** 10.1101/2025.10.29.680858

**Authors:** Simran Singh, Ashish Kumar, Sudipta Paul, Mriganka Sarkar, Santhosh Duraisamy, Harender Yadav, Seema Kuldeep, Soumita Bhaumik, Kunj Kumar Prajapati, Saahiba Thaleshwari, Anuj Gargya, Tamojit Santra, Sonal Amit, Rashmi Parihar, Santosh K. Misra, Hamim Zafar, Luiz F. Barella, Dharmaraja Allimuthu, Sai Prasad Pydi

**Author notes:** Corresponding author: Sai Prasad Pydi, PhD, Lab101, Mehta Family Centre for Engineering in Medicine Department of Biological Sciences & Bioengineering Indian Institute of Technology Kanpur, UP 208016, India. Equal contribution (1^st^ Author) (Simran Singh, Ashish Kumar). Equal contribution (2^nd^ Author) (Sudipta Paul, Mriganka Sarkar, Santhosh Duraisamy).

## Abstract

Obesity-associated inflammation disrupts pancreatic β-cell function, but the immune-derived signals that directly regulate insulin secretion remain incompletely defined. Here, we identify myeloid Gq signaling as a critical immunometabolic node that links macrophage activation to β-cell dysfunction. For the first time, we employed a chemogenetic approach (DREADDs) to selectively and temporally activate Gq-coupled GPCR signaling in myeloid cells to examine its effect on islet function. Our findings reveal that acute Gq activation in islet-resident macrophages impaired glucose-stimulated insulin secretion, uncovering a previously unrecognized immune–endocrine axis. Conversely, myeloid-specific Gαq deletion improves systemic glucose homeostasis, underscoring the physiological relevance of this pathway. Mechanistic analysis revealed that Gq activation in macrophages stimulates AMPK signaling and drives the secretion of sphingolipids. These lipids suppress insulin secretion and introduce a new mechanism for immune–islet communication, extending beyond traditional cytokine-based models. We further identify the lipid-sensing receptor GPR18 as an upstream activator of the Gq–AMPK pathway in macrophages. GPR18 stimulation recapitulated the Gq-dependent sphingolipid secretion and β-cell inhibitory phenotype, which was abolished in myeloid Gαq-deficient mice. Collectively, these findings establish a mechanistic framework whereby macrophage Gq signaling integrates lipid sensing and metabolic stress to modulate β-cell function. This work reveals a previously unrecognized macrophage–β-cell communication axis with therapeutic potential for restoring insulin secretion in metabolic diseases such as obesity and type 2 diabetes.

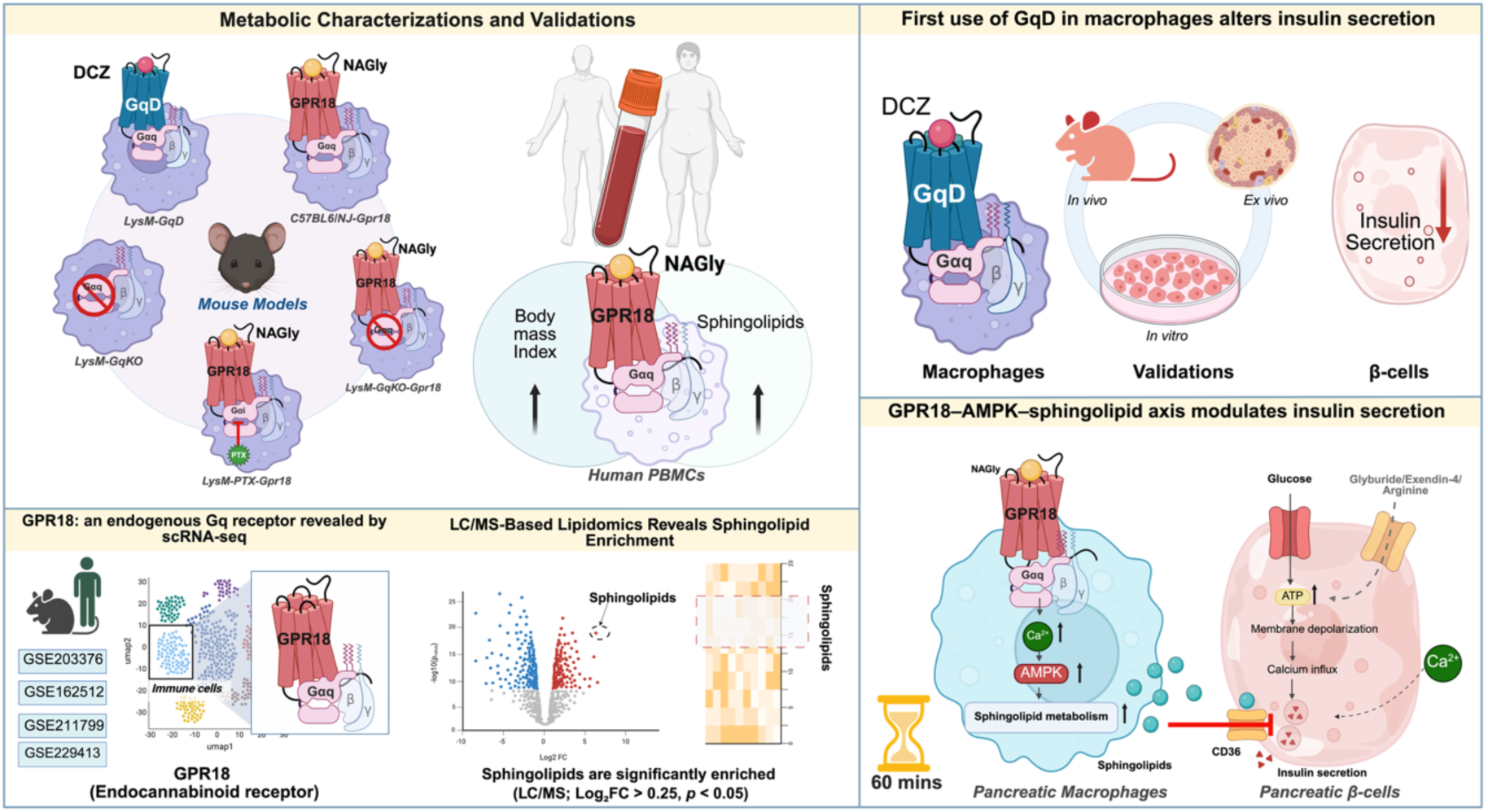

## Introduction

The pancreatic islet comprises a heterogeneous population of specialized cells functioning synergistically to carry out critical processes essential for insulin secretion and glucose homeostasis [1, 2]. In this microenvironment, β-cells play a central role in glucose-stimulated insulin secretion through a carefully orchestrated series of metabolic and signaling events. These include glucose sensing and uptake, glycolytic flux, mitochondrial oxidation, calcium influx, membrane depolarization, and insulin exocytosis [3, 4]. These steps can be compromised in the event of β-cell failure arising from various chronic metabolic conditions. Obesity is one of the major stressors for the β-cell function, where β-cells try to meet the insulin demand due to the concomitant development of insulin resistance (IR). Initially, β-cells adapt by augmenting insulin output; however, prolonged metabolic overload leads to β-cell exhaustion and failure. Besides IR, obesity can lead to low-grade inflammation, and the infiltration of immune cells significantly increases in various metabolic tissues, including pancreatic islets [5–7]. Evidence from multiple investigations indicates an increased infiltration of myeloid lineage cells, mainly monocytes and macrophages, into the pancreatic islets in both obese animal models and humans [8–11]. Intraislet macrophages exhibit significant functional plasticity; they impair β-cell function via direct cell-cell interaction and engulfment of insulin granules, and alternatively, they promote β-cell proliferation through platelet-derived growth factor receptor (PDGFR) signaling [6]. Emerging evidence suggests that blocking macrophage-derived exosomal miR-155 improves insulin sensitivity and glucose tolerance in obese and diabetic mice by enhancing β-cell function through the miR-155-PDX1 axis [12]. Following β-cell injury, islet-resident macrophages adopt a reparative phenotype, characterized by increased production of insulin-like growth factor 1 (IGF-1) and reduced expression of pro-inflammatory cytokines. This phenotypic shift supports insulin secretory capacity and contributes to metabolic homeostasis even under conditions of obesity and hyperglycemia [13]. These studies underscore the potential role of macrophages in obesity-associated β-cell dysfunction. However, the precise signaling pathways that govern macrophage activation and their interaction with β-cells remain to be elucidated. Identifying the molecular mediators involved in this intercellular crosstalk not only enhances our understanding of islet functioning but also unveils novel therapeutic targets.

Recent studies have shown that G protein-coupled receptors (GPCRs) are not only activated by classical ligands but also by various metabolic intermediates and secretory factors that play a central role in cellular metabolism. In line with their role in other tissues, few studies have explored the role of GPCRs in the pancreatic macrophages. Deficiency of GPR92 in islet macrophages can cause β-cell dysfunction, disrupting glucose homeostasis [14], and activation of FFAR4 on islet macrophages leads to IL6 release and promotes β-cell function [15]. Moreover, studies from murine models suggest that Ccl27a is expressed by pancreatic β-cells, where it interacts with its receptor Ccr2 present on islet macrophages; however, the signaling axis was disrupted upon prolonged high-fat diet exposure (Ccl27-Ccr2) [16]. Nevertheless, the function of macrophage GPCRs and the mechanisms governing their control of β-cell activity remain to be elucidated. This is primarily because many GPCRs are expressed across diverse cell types and tissues, making it challenging to delineate macrophage-specific effects. Further, the limited availability of highly selective agonists or antagonists for individual GPCRs has hindered mechanistic studies *in vivo*. Despite these challenges, comprehending the role of macrophage GPCRs in pancreatic islets is essential for advancing the development of innovative GPCR-targeted therapeutics for metabolic disorders. Specifically, understanding how acute and chronic activation of macrophage GPCR signaling affects β-cell function under physiological and pathological conditions would help develop better drugs for treating obesity and T2D.

Chemogenetics provides an effective means to circumvent the limitations associated with traditional receptor studies[17]. They also facilitate the expression of engineered receptors in a cell-specific manner, thereby allowing for precise temporal control of receptor activation. Previous studies have demonstrated that Gq signaling plays a diverse and crucial role in glucose and energy homeostasis within metabolically active tissues like pancreatic α- and β-cells, brown adipocytes, skeletal muscle and white adipocytes [18–22]. However, the role of myeloid cell Gq signaling, particularly under diet-induced obesity (DIO) models, is not fully characterized. To address this gap, in the current study, we expressed Gq DREADD in a cell-specific manner using the LyzM-Cre model and characterized its metabolic role in healthy and obese states. Interestingly, activation of Gq signaling in myeloid cells led to impaired insulin secretion from pancreatic β-cells, an effect mediated, at least in part, by sphingolipid species. Additionally, as proof of concept, we validated our findings using an endogenously expressed Gq-coupled receptor, GPR18, highlighting the relevance of this regulatory axis in both mice and human. Insights from this study will provide new guidance for developing better strategies to treat obesity and T2D.

## Results

### Macrophage Gnaq expression is selectively upregulated during obesity in humans and mice

To investigate whether Gαq G-protein expression is modulated in tissue-resident macrophages under metabolic stress, we integrated publicly available single-cell RNA sequencing (scRNA-seq) datasets generated from pancreatic islets of mice fed on regular chow (RC), high-fat diet (HFD), or diabetic db/db mice with 1,96,808 cells [23–26]. The integrated murine pancreatic atlas revealed eight distinct cell populations (**Figure 1A, S1A**), including one cell population classified as macrophages based on canonical marker expression (**Figure S1B**). Consistent with previous observations [16], macrophages were the predominant immune cell in RC and HFD conditions (**Figure 1B**). In contrast, db/db islets exhibited a markedly distinct immune cell composition (**Figure S1C**), indicating a diet-independent, immune remodelling exacerbating β-cell dysfunction. Our prior scRNA-seq study [16] demonstrated substantial heterogeneity within the islet macrophage compartment, with a predominance of pro-inflammatory (M1-like) phenotype. Hence, we specifically examined *Gnaq* expression within this macrophage cell type. Interestingly, *Gnaq* expression levels were elevated in M1macrophages from HFD islets, suggesting a diet-dependent alteration in Gq signaling (**Figure 1C**). Given the functional overlap between Gq and G11 proteins, we next assessed *Gna11* expression in myeloid cells and interestingly, *Gna11* expression remained unchanged between RC and HFD conditions (**Figure 1C**).

**Figure 1.**
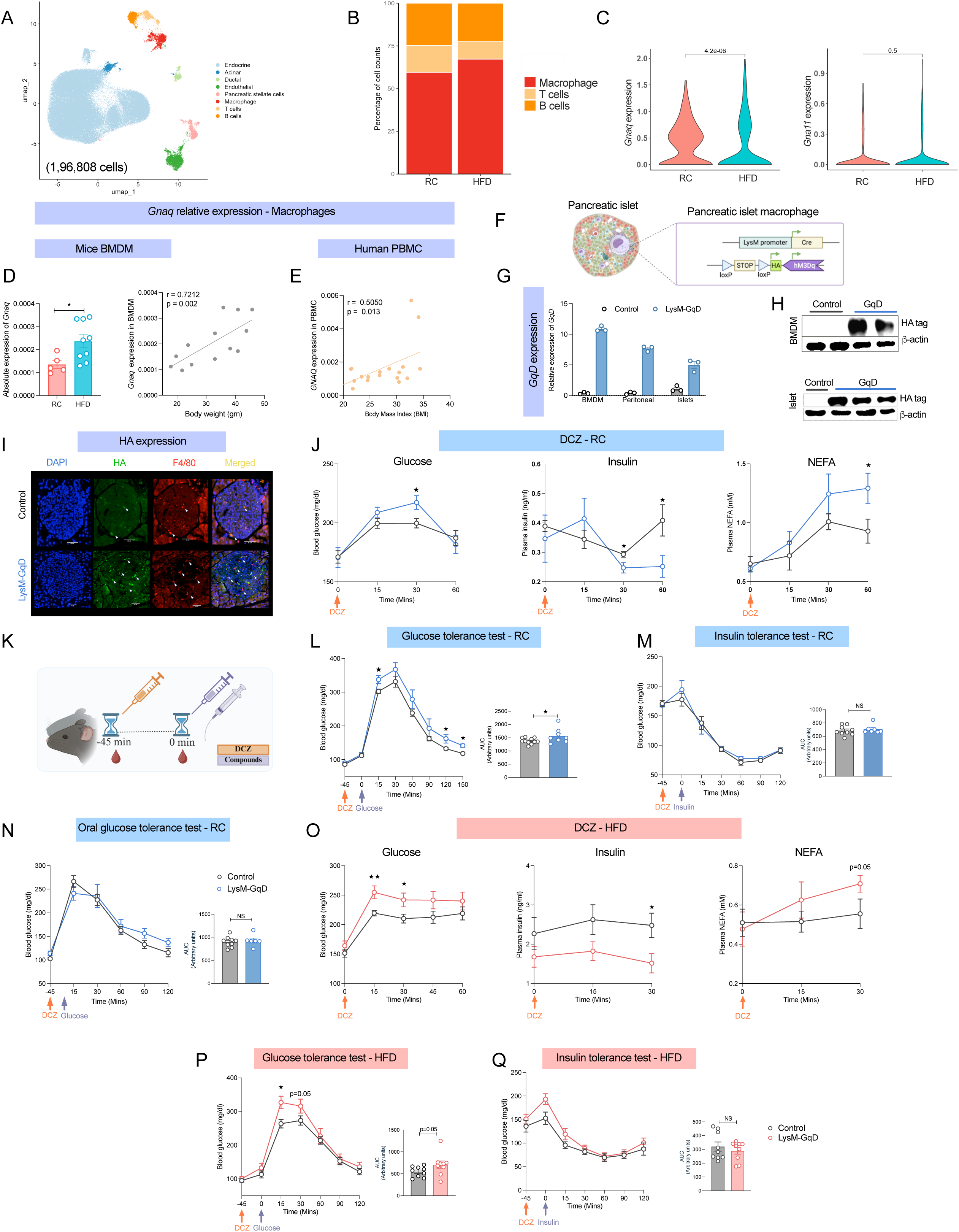
Acute activation of myeloid Gq signaling perturbs glucose homeostasis in lean and obese mice. (A) UMAP showing murine pancreatic islet cell types from mice fed regular chow (RC), high-fat diet (HFD), or with a db/db background (1,96,808 cells). (B) Distribution of pancreatic islet immune cell types in scRNA seq RC- and HFD-fed mice. (C) Expression levels of *Gnaq* (left) and *Gna11* (right) in scRNA seq pancreatic islet macrophages maintained on RC and HFD conditions. (D) Quantitative PCR indicating *Gnaq* expression in murine bone marrow– derived macrophages (BMDMs) (left) and the correlation of *Gnaq* expression with body weight in BMDMs from RC (n=5) and HFD (n=9) mice (right) (E) Correlation of human peripheral blood mononuclear cells (PBMCs) *GNAQ* expression and with body mass index (BMI) (n=19) (E). (F) Schematic representation of the LysM-GqD construct with floxed Gq-DREADD under the lysozyme-2 promoter driving Cre in pancreatic islet macrophages. (G) GqD expression levels in BMDMs, peritoneal macrophages, and pancreatic islets from control and LysM-GqD mice (n=3). (H) Immunoblot confirming HA-tagged GqD expression in BMDMs (top) and pancreatic islets (bottom), with β-actin as a loading control (n=2–3). (I) Immunofluorescence of pancreatic islets showing HA-tagged GqD in F4/80⁺ macrophages (DAPI, blue; F4/80, red; HA, green), scale-20µM (n=3). (J) Blood glucose (left), plasma insulin (middle), and plasma non-esterified free fatty acid (NEFA) (right) levels in RC-fed LysM-GqD and control mice following acute DCZ administration (30 µg/kg, i.p.; n=6–9). (K) Schematic illustration of the metabolic testing workflow outlining DCZ activation and compound administration procedure. (L) Glucose tolerance test (GTT; 2 g/kg glucose, i.p.; n=8–11) (M) Insulin tolerance test (ITT; 0.75 IU/kg insulin, i.p.; n=7–9) (N) Oral glucose tolerance test (OGTT; 2 g/kg glucose via gavage; n=6–9) in RC-fed LysM-GqD and control mice following acute DCZ administration (30 µg/kg, i.p.). (O) Blood glucose (left), plasma insulin (middle), and plasma NEFA (right) levels in HFD-fed LysM-GqD and control mice following acute DCZ administration (30 µg/kg, i.p.; n = 6–10). (P) GTT (1 g/kg glucose, i.p.; n=9), (Q) ITT in HFD-fed mice (1 IU/kg insulin, i.p.; n=9) with corresponding AUC analysis (right) (n=9) in HFD-fed LysM-GqD and control mice following DCZ administration (30 µg/kg, i.p.). Data are shown as mean ± s.e.m. using one-tailed unpaired t-test: p < 0.05, *p < 0.01, NS = not significant (D, E, G, J, L–Q).

To experimentally validate these findings, we assessed *Gnaq* expression in bone marrow–derived macrophages (BMDMs) from RC and HFD mice. Consistent with the scRNA-seq findings, *Gnaq* expression was significantly upregulated under HFD and positively correlated with body weight (**Figure 1D**). In contrast, *Gna11* expression in BMDMs exhibited no significant difference or correlation with body weight across dietary conditions (**Figures S1D and E**). However, in peritoneal cells, we found that both *Gnaq* and *Gna11* were upregulated in HFD intervention and positively correlated to body weight and blood glucose as compared to the RC cohort (**Figure S1F-J**).

To evaluate the translational relevance of these findings, we quantified *GNAQ* expression in human peripheral blood mononuclear cells (PBMCs) and observed a positive correlation with body mass index (BMI) (**Figure 1E**). However, no significant correlation was found between *GNAQ* mRNA levels and fasting glucose in human PBMCs (**Figure S1K**). Similarly, *GNA11* expression showed no significant correlation with BMI or fasting glucose in the human samples (**Figures S1L and M**). Collectively, these results indicate that *Gnaq* protein expression in macrophages is dynamically regulated in obesity in both mice and humans. The selective increase of *Gnaq* in macrophages indicates a possible functional consequence in metabolic stress conditions.

### Acute activation of myeloid Gq signaling disrupts glucose homeostasis in lean and obese mice

To evaluate the functional impact of myeloid Gq signaling on systemic metabolism, we generated mice expressing a Gq-coupled DREADD receptor under the control of LysM-Cre (LysM-GqD), targeting the myeloid lineage. From here on, we refer to Cre-positive mice as LysM-GqD and Cre-negative littermates carrying only the floxed allele as control (**Figure 1F**). Transgene expression was confirmed by qPCR in BMDMs, peritoneal cells and islets (**Figure 1G**), and by immunoblotting and immunofluorescence using an HA-tag (**Figures 1H and I**), confirming selective expression in islet macrophages using F4/80^+^ Ab. To determine whether the Gq-DREADD receptor displayed any constitutive activity, we compared body weights, glucose tolerance, and glucose-stimulated insulin secretion (GSIS) between control and LysM-GqD mice in the absence of DCZ, the synthetic ligand for DREADD. No significant differences were observed in these baseline parameters (**Figures S2A-C**), indicating that the receptor remains inactive without ligand stimulation. We first evaluated the systemic metabolic impact of acute myeloid Gq activation without assuming tissue specificity due to the widespread distribution of macrophages throughout metabolically active tissues. Upon acute DCZ administration, LysM-GqD mice exhibited elevated blood glucose, reduced plasma insulin at 30 minutes, and increased NEFA levels at 60 minutes (**Figure 1J**). As we observed metabolic alterations at least 30 minutes after administration and considering that macrophages are not direct regulators of systemic glycemia, we hypothesized that macrophage-derived factors may influence glucose homeostasis indirectly. Therefore, all subsequent metabolic assays were conducted 45 minutes following DCZ injection (**Figure 1K**). We tested two different doses of DCZ (10 µg/kg and 30 µg/kg). Both concentrations produced similar phenotypes; however, the effects were more pronounced at the higher dose (30 µg/kg) (**Figure S2D**). Therefore, we selected 30 µg/kg as the working dose for all subsequent experiments. Under RC conditions, activation of Gq signaling in myeloid cells impaired glucose and pyruvate tolerance, suggesting altered glucose clearance (**Figure 1L and S2E**). However, insulin tolerance tests (ITT) revealed no significant differences between groups, indicating that insulin sensitivity remained intact (**Figure 1M**). Similarly, oral glucose tolerance testing (OGTT) did not show significant changes, suggesting that enteral glucose sensing and absorption were not significantly affected (**Figure 1N**).

Next, to evaluate whether these effects were maintained in the context of diet-induced obesity, we maintained control and LysM-GqD mice on an HFD for at least 12 weeks prior to experimentation. Despite no difference in body weight between groups (**Figure S2F**), Gq activation under HFD settings resulted in a metabolic profile comparable to that of lean mice, including higher blood glucose levels, decreased insulin secretion, and elevated plasma FFA (**Figure 1O**). Further, glucose and pyruvate tolerance were compromised (**Figure 2P and S2G**); however, insulin tolerance and oral glucose tolerance were unaffected (**Figure 1Q and S2H**). These findings suggest that myeloid Gq signaling primarily influences systemic glucose metabolism through impaired glucose clearance, rather than altering insulin sensitivity. We also evaluated the metabolic effects like energy expenditure and respiratory exchange ratio (RER) upon acute myeloid Gq signaling in both RC and HFD intervention. Interestingly, we did not find any significant difference in either healthy or obese state, suggesting effects are limited to impaired glucose clearance (**Figure S3A-P**).

**Figure 2.**
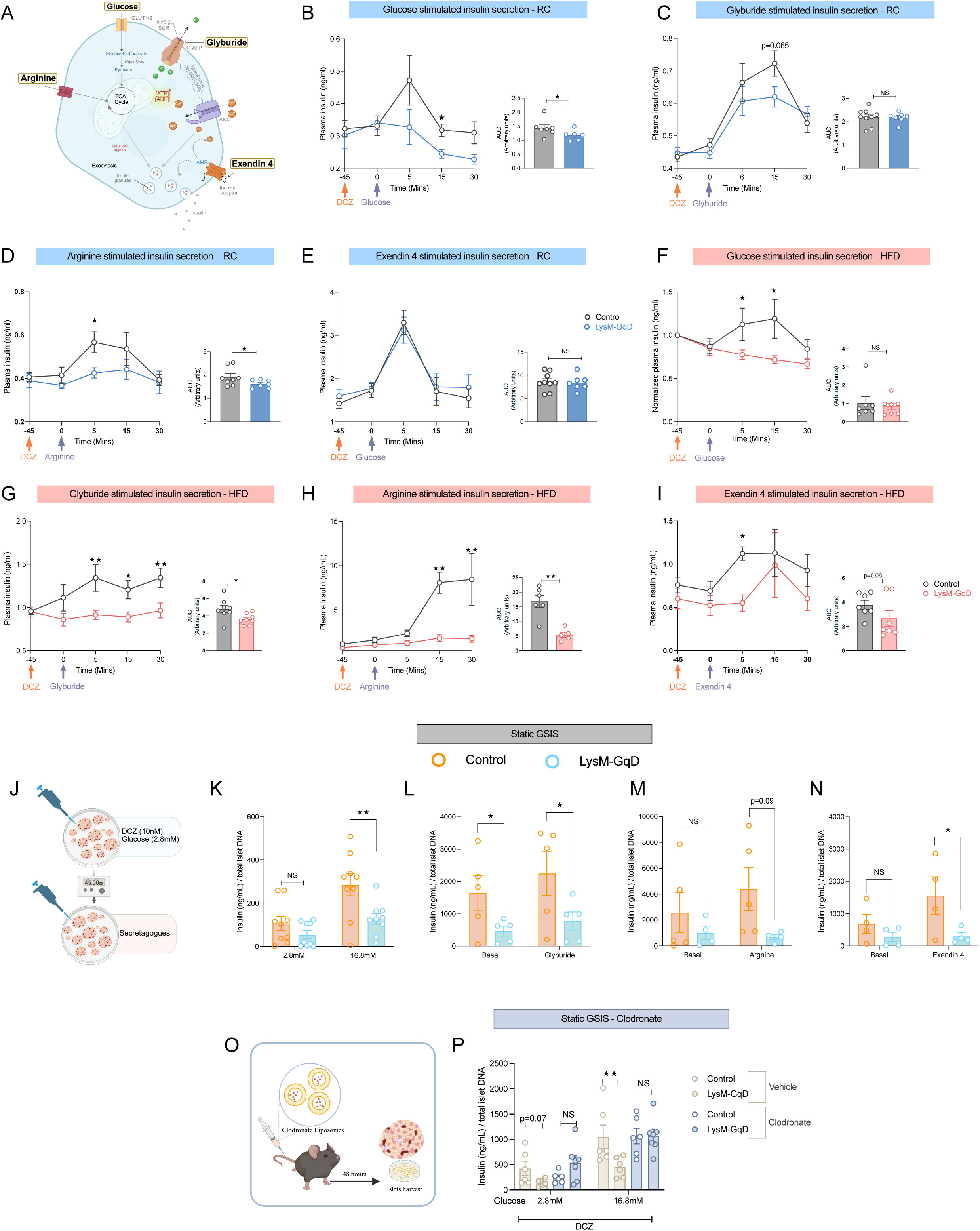
Myeloid-specific GqD activation impairs insulin secretion *in vivo* and *ex vivo*. (A) Schematic representing of pharmacological agents used to stimulate insulin secretion in β-cells. (B–E) Plasma insulin levels in RC-fed control and LysM-GqD mice following acute DCZ administration (30 µg/kg, i.p., 45 min) in response to glucose (2 g/kg, i.p.; n = 6–7) (B), glyburide (5 mg/kg, i.p.; n = 7–8) (C), L-arginine (1 g/kg, i.p.; n = 7–8) (D), or exendin-4 (12 nmol/kg, i.p.; n = 7–10) (E). (F–I) Plasma insulin levels in HFD-fed control and LysM-GqD mice after acute DCZ stimulation in response to glucose (1 g/kg, i.p.; n = 8) (F), glyburide (10 mg/kg, i.p.; n = 8) (G), L-arginine (1 g/kg, i.p.; n = 5–6) (H), or exendin-4 (4 nmol/kg, i.p.; n = 7) (I), with corresponding AUC analyses (right). (J) Schematic representation of the *ex vivo* insulin secretion assay in isolated pancreatic islets. (K–N) Static insulin secretion from isolated islets treated with DCZ (30 nM) under low (2.8 mM) and high (16.8 mM) glucose conditions (n = 9) (K), glyburide (100 nM; n = 5) (L), L-arginine (30 mM; n = 4–5) (M), or exendin-4 (n = 4) (N), normalized to total islet DNA content. (O) Schematic representation of clodronate liposome treatment for macrophage depletion in *ex vivo* islet studies. (P) Static glucose-stimulated insulin secretion (GSIS) in macrophage-depleted islets from control and LysM-GqD mice following clodronate treatment (10 µL/g, i.v.; n = 5). Data are presented as mean ± s.e.m. Statistical tests: two-tailed unpaired *t*-test (B–E, AUC analyses), two-way ANOVA (F–I), one-tailed unpaired *t*-test (K–N, P). *p* < 0.05, \**p* < 0.01, \*\**p* < 0.001, NS = not significant.

### Myeloid Gq signaling acute activation suppresses β-cell insulin secretion in both lean and obese mice

To test whether the impaired glucose tolerance observed in myeloid Gq is due to changes in insulin secretion, we evaluated β-cell response to different insulin secretagogues under RC and HFD conditions (**Figure 2A**). First, we administered glucose to the LysM-GqD and control groups maintained on RC diet and measured plasma insulin levels at regular intervals. Interestingly, plasma insulin was significantly reduced in the experimental group, indicating a defect in GSIS (**Figure 2B**). Next, we tested glyburide and arginine to determine whether the suppression of insulin secretion is specific to glucose or to other insulinotropic stimuli. These secretagogues bypass glucose metabolism and stimulate insulin release through direct β-cell depolarization. In response to glyburide, LysM-GqD mice showed a drop in insulin levels 15 minutes after injection (**Figure 2C**). Additionally, arginine causes a weaker insulin response than controls, indicating that activating Gq in myeloid cells generally hampers the β-cells ability to secrete insulin (**Figure 2D**). Surprisingly, insulin secretion in response to exendin-4, a GLP-1 receptor agonist that enhances insulin secretion via cAMP-dependent pathways independently of glucose transport, was not altered between LysM-GqD and control group mice under RC conditions (**Figure 2E**).

In HFD fed mice, activating myeloid Gq similarly reduced insulin secretion when responding to glucose, glyburide, and arginine, indicating a broad β-cell secretory impairment (**Figure 2F-H**). Remarkably, insulin secretion stimulated by exendin-4 was also decreased, suggesting that obesity increases β-cells sensitivity to myeloid Gq- related suppression, even with GLP-1 receptor activation (**Figure 2I**). These findings demonstrate that acute activation of myeloid Gq signaling reduces β-cell insulin secretion across various stimuli, and HFD exposure further exacerbates this dysfunction. We believe these effects might result from disrupted immune-islet communication that disrupts β-cell function and impaired glucose homeostasis.

As we observed impaired insulin secretion *in vivo*, to determine whether the myeloid cells in the islet microenvironment are sufficient to produce these effects, we isolated the islets from the LysM-GqD and control mice, and treated them with various insulin secretagogues in the presence of DCZ (**Figure 2J**). In agreement with our hypothesis, DCZ-induced activation of Gq signaling in myeloid cells significantly decreased insulin secretion from isolated islets in response to high glucose, glyburide, exendin-4, and arginine (**Figure 2K-N**). These results show that Gq-induced impairments are not restricted to systemic effects but can also result from local macrophage–islet interactions.

Next, to confirm that the effects arise from the intra-islet macrophages, we treated both LysM-GqD and control mice with clodronate and vehicle liposomes (**Figure 2O**). Macrophage depletion efficiency was confirmed by immunofluorescence and qPCR analysis (**Figure S4A-C**). In LysM-GqD islets, insulin secretion under high glucose stimulation was significantly impaired; however, clodronate-mediated macrophage depletion restored insulin secretory responses (**Figure 2P**), implicating islet macrophages as mediators of the β-cell dysfunction observed upon Gq activation.

### Myeloid Gq activation impairs β-cell function via macrophage-derived paracrine lipid signals

Since pancreatic islets are composed of multiple cell types, to further dissect the direct macrophage–β-cell crosstalk, we employed a co-culture system using BMDMs and INS- 1 β-cells at a 1:5 ratio, reflecting the relative abundance of macrophages to β-cells in islets from HFD-fed mice [6] (**Figure 3A**). Conditioned media from DCZ-stimulated LysM- GqD BMDMs robustly suppressed insulin secretion from INS-1 cells in response to high glucose, glyburide, exendin-4, and arginine, mirroring the *in vivo* and *ex vivo* findings (**Figure 3B-C**). In contrast, stimulation with FPL64176, a voltage-gated calcium channel activator that directly bypasses upstream signaling to trigger insulin granule exocytosis, elicited comparable insulin secretion between groups (**Figure 3C**). These results suggest that β-cell calcium-dependent secretory machinery remains intact and that Gq-induced macrophage signals act upstream of calcium entry.

**Figure 3.**
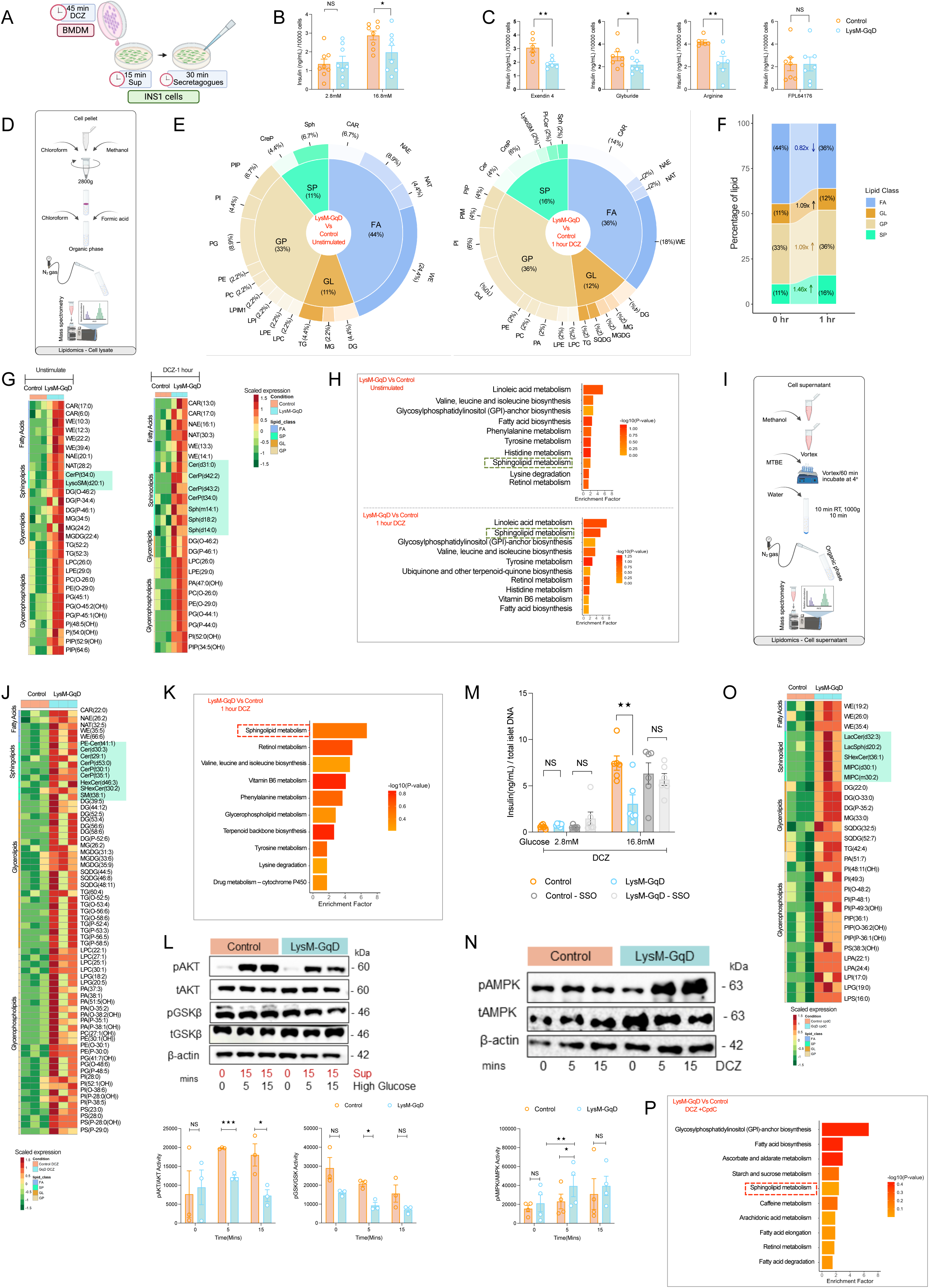
Myeloid Gq activation impairs β-cell function via macrophage-derived paracrine lipid signals. (A) Schematic representation of the indirect co-culture strategy in which supernatants from BMDMs stimulated with DCZ (45 min) were transferred to INS-1 β-cells. (B– C) Insulin secretion from INS-1 cells cocultured with control or LysM-GqD BMDM supernatants in response to DCZ (30 nM) under low (2.8 mM) and high (16.8 mM) glucose (n = 9) (B), or under exendin-4 (12 nM; n = 4), glyburide (100 nM; n = 5), L-arginine (30 mM; n = 4–5), or FPL64176 (0.1 µM; n=7)(C). (D) Workflow for lipid extraction from BMDM cell pellets for untargeted lipidomics. (E) Differentially abundant lipid species in control and LysM-GqD BMDMs under basal conditions and after 1 hour DCZ stimulation (n = 3). (F) Proportional increase in lipid species after 1 hour GqD activation with DCZ compared to unstimulated BMDMs (n = 3). (G) Heatmaps of lipid species in untreated (left) and DCZ-treated (1 hour, right) BMDMs (n = 3) from control and LysM-GqD mice. (H) Pathway enrichment analysis depicting upregulated pathways in LysM-GqD BMDMs compared to control BMDMs under untreated conditions (top) and following 1 hour DCZ treatment (bottom; n = 3). (I) Workflow for lipid extraction from BMDM supernatants for untargeted lipidomics. (J) Heatmaps showing lipid species in the supernatant of BMDMs from control and LysM-GqD mice after 1 hour DCZ treatment (n = 3) (K) Pathway enrichment analysis showing upregulated pathways in the supernatant of LysM-GqD BMDMs compared to control BMDMs following 1 hour DCZ treatment (n = 3). (L) Immunoblot of INS-1 cells showing AKT and GSK-β phosphorylation after exposure to supernatants from DCZ-stimulated BMDMs under high-glucose (16.8 mM) conditions for 5 and 15 min (n = 3). (M) Insulin secretion in INS-1 cells after transfer of supernatants from control or LysM-GqD BMDMs stimulated with DCZ (30 nM) at 2.8 or 16.8 mM glucose, with or without the CD36 inhibitor sulfosuccinimidyl-oleate (SSO; 0.5 mmol/l). (N) Immunoblot showing AMPK phosphorylation in BMDMs after acute GqD activation with DCZ (0, 5, and 15 min). (O) Heatmaps showing lipid species in BMDMs from control or LysM-GqD BMDMs treated with the ampk inhibitor compound C (10 µM) during DCZ stimulation (n = 3). (P) Pathway enrichment analysis showing enriched pathways in BMDMs treated with compound C during DCZ stimulation from control or LysM-GqD (n = 3). Data are presented as mean ± s.e.m. One-tailed unpaired t-test (B–C); two-tailed unpaired t-test (N–O). *p < 0.05, **p < 0.01, ***p < 0.001, NS = not significant. SP, sphingolipids; FA, fatty acids; GP, glycerophospholipids; GL, glycerolipids.

Given the role of Gq signaling in lipid homeostasis, we sought to determine how lipid dysregulation contributes β-cell dysfunctions. Therefore, we performed an untargeted lipidomic analysis to investigate whether any secretory factors released by macrophages contribute to β-cell dysfunction. To test this hypothesis, we stimulated BMDMs from LysM-GqD and control mice with DCZ for 0, 60, and 120 minutes (**Figure 3D**). Interestingly, we observed accumulation of sphingolipid species in DCZ-treated cells, with more pronounced levels in both 60 and 120 min. treatment. The PCA analysis of untreated BMDM from control and LysM-GqD did not alter, however upon DCZ treatment for 60 and 120 min revealed altered lipidome (**Figure S5A**). We next analysed the differentially regulated lipids across 0, 60 and 120 min and found the different species of sphingolipids were upregulated upon DCZ-GqD activation in 60 and 120 min when compared to untreated cells (**Figure S5B**). These sub class of lipids species majorly belong to sphingolipid (SP) class, diversity of the sphingolipid subtypes was highly diverse in DCZ-GqD activation in 60 min and 120 min of stimulation and increases by ∼ 1.46 folds in both 60 and 120 min as compared to 0 mins (**Figure 3E and F, S5C and D**). However, such diversity was not uniquely present in any of other lipid classes including, fatty acids (FA), glycerolipids (GL) and glycerophospholipids (GP) (**Figure 3G and S5E**). Further, enrichment analysis also pointed out highly enriched metabolic pathways related to sphingolipid metabolism within 60 min and 120 min of activation (**Figure 3H and S5F**). We next asked whether these lipids were secreted and available to act as a paracrine signaling molecule. To test this, we isolated lipids from the cell culture supernatants of DCZ-treated macrophages (**Figure 3I**). The partial least squares discriminant analyses (PLSDA) of control and LysM-GqD stimulated with DCZ substantially altered the BMDM lipidome (**Figure S5G**). Several sphingolipid species, including different subtypes of ceramides, were significantly enriched in extracellular media from the LysM-GqD group, indicating active secretion accompanied with highly enriched sphingolipid metabolic pathway (**Figure 3J and K**, **S5H-I**).

Since sphingolipids can interfere with insulin signaling, we examined their functional effects on β-cells. INS-1 cells were cultured in conditioned media from either control or LysM-GqD macrophages, and signaling was assessed following glucose stimulation. Cells exposed to LysM-GqD-derived supernatants showed markedly reduced phosphorylation of several insulin signaling components, including AKT, and GSK3β (**Figure 3L**), indicating compromised intracellular insulin signaling in β-cells. CD36 is known to transport lipid species such as ceramides into cells [27]. Next, we tested whether blocking this transporter could prevent signaling disruption. INS-1 cells were pretreated with a CD36 inhibitor (SSO) before exposure to conditioned media from LysM-GqD macrophages. Strikingly, this intervention fully restored insulin pathway activation, indicating that the observed effects were CD36-dependent (**Figure 3M**). Overall, these findings support a model in which myeloid Gq signaling promotes the release of bioactive sphingolipids that impair β-cell insulin signaling and alter insulin secretion.

### Macrophage-derived sphingolipid secretion is mediated by the AMPK signaling axis downstream of Gq activation

As we know, activation of Gq signaling in macrophages induces sphingolipid secretion and leads to β-cell dysfunction, next we sought to delineate the molecular mechanism underlying this lipid release. Given the role of AMPK in lipid metabolism [28], we examined the activation status following acute Gq activation. As expected, upon Gq activation with DCZ, phosphorylation of AMPK increased within 5 to 15 min (**Figure 3N**). To determine whether AMPK can contribute to the observed phenotype, we treated LysM-GqD BMDM with AMPK pharmacological inhibitor Compound C (CpdC), followed by DCZ stimulation for 60 min. PLSDA of control and LysM-GqD stimulated with CpdC and DCZ substantially altered the BMDM lipidome (**Figure S5I**). Lipidomic analysis of cell-free supernatants revealed that there was an alteration in the subspecies of sphingolipids as compared to without inhibitor pre-treatment. Additionally, the sphingolipid metabolism was not highly enriched (**Figure 3O and P**). These results indicate that the Gq–AMPK axis acts as an early signaling cascade in orchestrating the release of bioactive lipids that impair β-cell function.

### Myeloid-specific deletion of Gαq leads to improved glucose clearance

Since activation of macrophage Gq signaling impairs insulin secretion, we next investigated whether impaired Gq signaling would influence insulin secretion. To evaluate our hypothesis, we generated LysM-Cre–mediated *Gnaq* knockout mice (LysM-GqKO) (**Figure 4A**) and performed a series of metabolic tests. Throughout the study, no significant difference in body weight was observed between the LysM-GqKO mice and their littermate controls (**Figure 4B**) in healthy conditions. Fasting blood glucose levels were similar between the groups; however, fed blood glucose levels were significantly lower in LysM-Gq KO mice (**Figure 4C**), indicating improved glycaemic regulation. But, no significant difference in plasma insulin was observed between the groups, in the fasting and fed state (**Figure 4D**). Next, we performed GTT by administering glucose i.p., and LysM-Gq KO mice displayed improved glucose clearance (**Figure S6A**). In contrast, no significant difference in blood glucose levels was observed between the groups for ITT , PTT and OGTT (**Figure S6B-D**).

**Figure 4.**
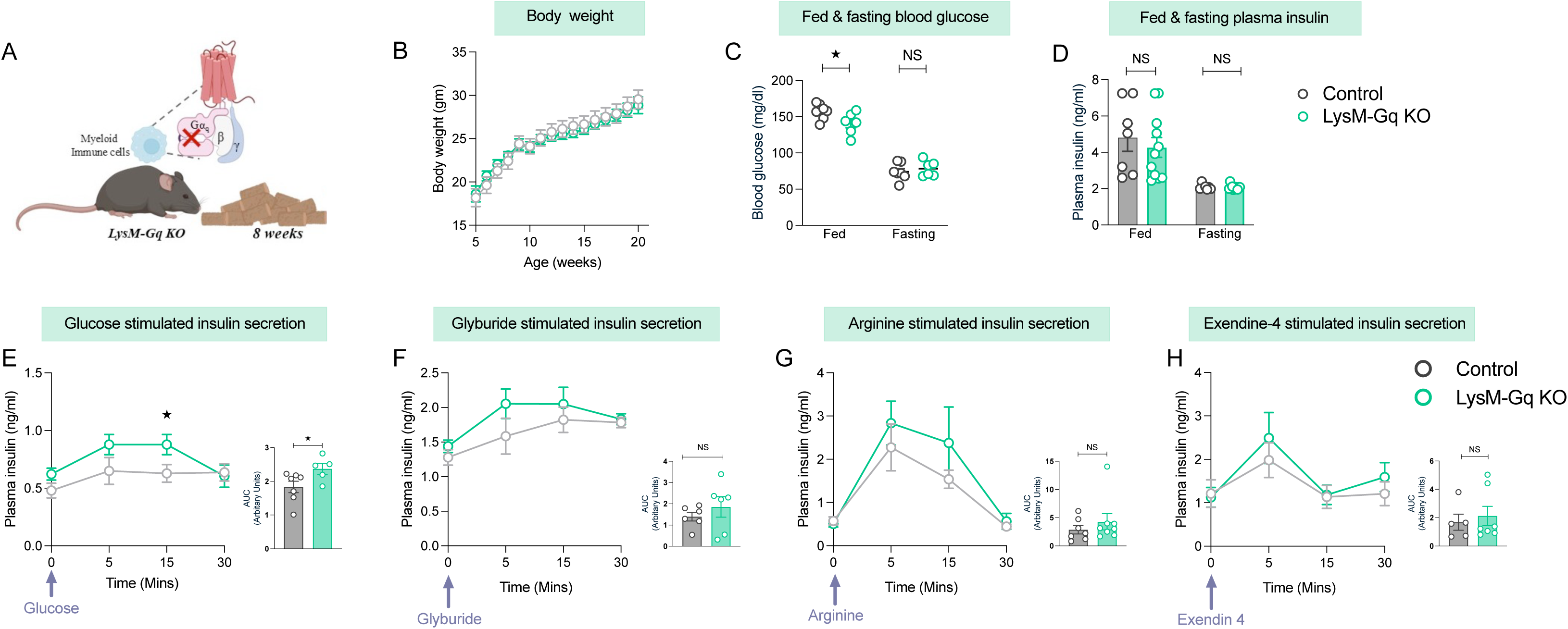
Metabolic characterization of LysM-Gq knockout mice shows improved glucose tolerance. (A) Schematic representation of the LysM-Gq KO model. (B) Body weight of control and LysM-Gq KO littermates was measured weekly from 4–21 weeks of age (n=6-7), maintained on RC diet. (C) Random-fed and fasting blood glucose levels in control and LysM-Gq KO mice (n=6-7). (D) Plasma insulin levels in fed and fasting states of control and LysM-Gq KO littermates maintained on RC diet (n=7-10). (E–H) Insulin secretion profiles in response to metabolic secretagogues in control and LysM-Gq KO maintained on RC diet: glucose (2 g/kg, i.p.) (n=5-7) (E), glyburide (10 mg/kg, i.p.) (n=6) (F), L-arginine (1 g/kg, i.p.) (n=7-8) (G), and exendin 4 (12 nmol/kg, i.p.) (n=5-9) (H). Line graphs show plasma insulin levels at indicated time points, with corresponding AUC analyses shown to the right. Data are presented as mean ± s.e.m. One-tailed unpaired t-test (C-H) *p < 0.05, **p < 0.01, ***p < 0.001; NS = not significant.

To determine whether enhanced glucose tolerance in LysM-GqKO mice was related to β-cell function, we assessed GSIS. After glucose administration, LysM-GqKO mice showed a significant increase in insulin secretion at 5 minutes compared to controls (Figure 4E), indicating a trend towards improved β-cell secretory capacity. In contrast, there was no significant difference between groups in insulin secretion in response to glyburide, arginine and exendin-4 (**Figures 4F-H**). These findings collectively show that loss of Gαq signaling in myeloid cells enhances systemic glucose clearance and boosts β-cell insulin secretion, while not changing insulin sensitivity or impacting other plasma metabolites like glycerol, triglycerides and free fatty acids (**Figure S6E-G**). We also confirmed the *Gnaq* expression in both control and LysM-Gq KO mice, we found that expression of *Gnaq* was reduced in knock out model (**Figure S6H**).

### GPR18 activation in macrophages suppresses insulin secretion via Gαq-dependent signaling

Given that myeloid-specific activation of Gq inhibits insulin secretion, we next asked whether activation of an endogenous Gαq-coupled GPCR in macrophages could recapitulate the inhibitory phenotype observed in LysM-GqD mice. To identify candidate receptors, we interrogated publicly available scRNA-seq datasets of murine macrophages (data not shown) and compiled the top ten GPCRs expressed across subsets. We then performed quantitative PCR validation in primary bone marrow–derived macrophages. Notably, *GPR18* was consistently and significantly enriched in macrophages, with expression further elevated in classically activated M1 macrophages relative to M0 controls (**Figure S7A**).

Next, to confirm its expression in islet-resident macrophages, we analyzed islet scRNA-seq datasets, and *Gpr18* was expressed in immune clusters (**Figure 5A and S7B**). To confirm the cell-specific expression of *Gpr18*, we quantified *Gpr18* mRNA expression levels in BMDMs, isolated islets, and INS-1 β-cell lines. In agreement with the scRNA-seq data, *Gpr18* was expressed in BMDMs and whole islets but was virtually undetectable in INS-1 cells (**Figure 5B**). To further investigate the expression pattern of Gpr18 in macrophages, we examined its expression under different conditions. We found that *Gpr18* was upregulated in HFD peritoneal cells (**Figure 5C**) and positively correlated with body weight and blood glucose (**Figure 5D**). However, in BMDMs from HFD mice, *Gpr18* expression was downregulated (**Figure S7C-F**), contrasting with our observations in freshly isolated peritoneal cells from HFD mice. Together, these findings reveal that both metabolic states and inflammatory polarization dynamically regulate *Gpr18* expression in macrophages. As GPCRs can exhibit context-dependent G protein coupling, we confirmed that *GPR18* activation engages Gq signaling in macrophages. Using a homogenous time-resolved fluorescence (HTRF)-based IP1 assay, stimulation of BMDMs with the known GPR18 agonist *N*-arachidonoyl glycine (NAGly) induced a dose-dependent accumulation of IP1, consistent with canonical Gq signaling (**Figure S7F**).

**Figure 5.**
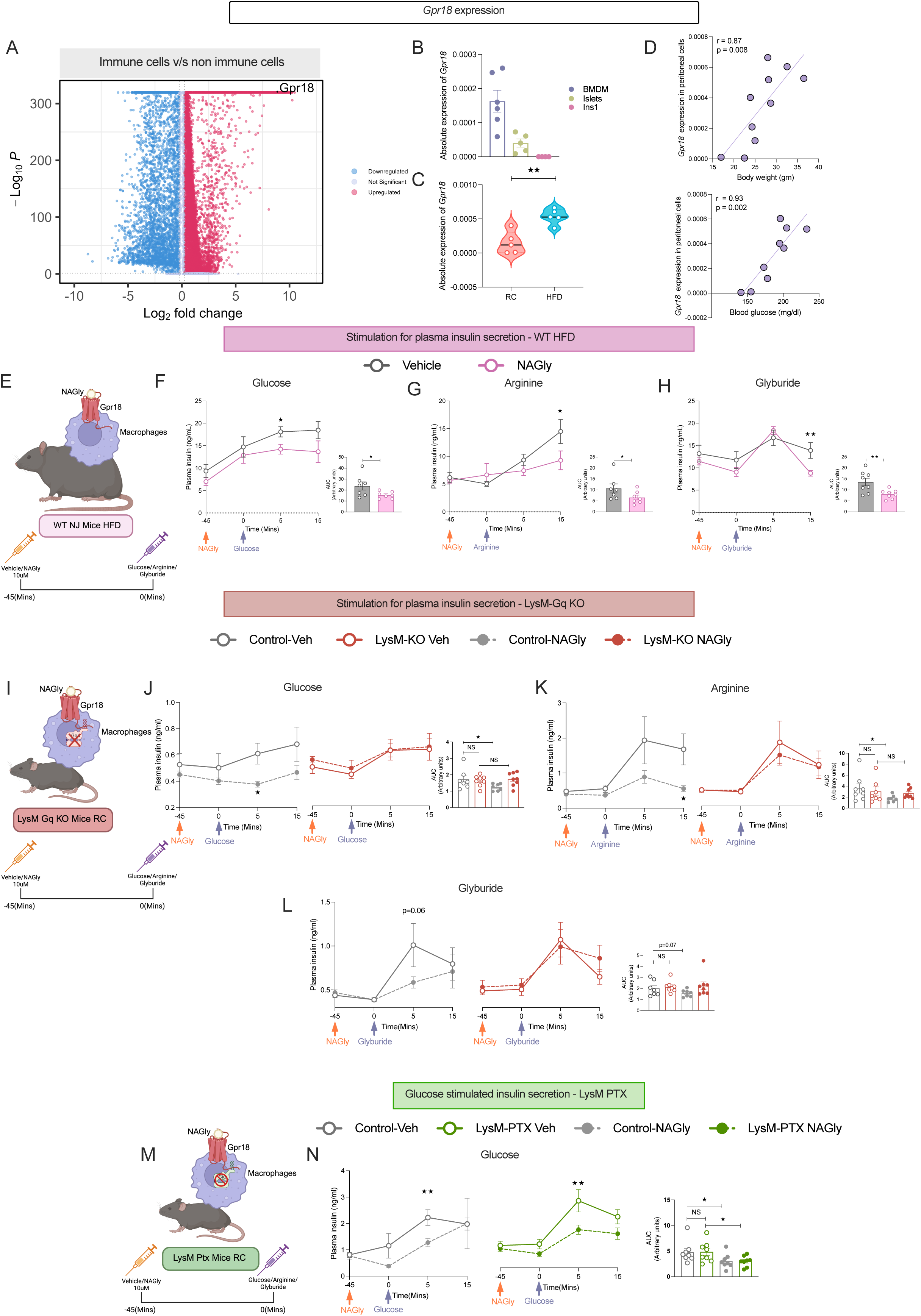
GPR18 activation in macrophages suppresses insulin secretion via Gq-dependent signaling. (A) Volcano plot showing upregulated expression of *Gpr18* in scRNA seq pancreatic islet immune cells from RC, HFD fed and db/db background mice. (B, C) Relative *Gpr18* expression in BMDMs, islets, INS-1 cells (B), and peritoneal macrophages of mice fed on RC or HFD (C). (D) The correlation of the Gpr18 expression with body weights and blood glucose levels in mice fed on RC and HFD. (E) Schematic representation of experimental design in WT HFD-fed mice treated with NAGly to activate GPR18. (F-H) Plasma insulin secretion in WT HFD-fed mice treated with vehicle or NAGly (36.15 µg/kg) in response to glucose (n=7) (F), L-arginine (n=8) (G), or glyburide (n=8) (H); corresponding AUC analyses shown. Schematic of experimental design in LysM-Gq KO mice under RC diet. (J-L). Plasma insulin secretion in control (left) and LysM-Gq KO (right) mice treated with vehicle or NAGly in response to glucose (n=8) (J), L-arginine (n=8) (K), or glyburide (n=8) (L). (M) Schematic representation of experimental design in LysM-PTX KO mice. (N) Plasma insulin secretion in control (left) and LysM-PTX (right) KO mice in response to glucose with or without NAGly treatment (n=8). Line graphs show plasma insulin levels at the indicated time points, with corresponding AUC analyses shown to the right Data are presented as mean ± s.e.m. using one-tailed unpaired t-test (C-H) *p < 0.05, **p < 0.01, ***p < 0.001; NS; not significant.

To test the functional relevance of this pathway *in vivo*, we administered 36.16µg/kg of NAGly to C57BL/6J mice maintained on HFD. No significant difference in weight gain was observed between vehicle- and NAGly-treated cohorts (**Figure S7G**). 45 minutes post-NAGly injection, mice were challenged with either glucose, arginine, or glyburide, and plasma insulin levels were measured at baseline and at 5 and 15 minutes post-stimulation (**Figure 5E**). NAGly-treated animals significantly reduced insulin secretion in response to glucose (5 min), arginine (15 min), and glyburide (15 min) compared to vehicle controls. The AUC was significantly different in case of glucose and glyburide but not in arginine (**Figure 5F-H**). It is well known that GPCRs are expressed in multiple cell and tissue types, hence, to determine whether these effects were mediated through myeloid Gq signaling, we performed identical pharmacological experiments in LysM-Gq KO *and* control mice (**Figure 5I**). Interestingly, NAGly administration failed to suppress insulin secretion in the knockout animals, implicating myeloid Gq as an essential component of *Gpr18*-mediated β-cell inhibition (**Figure 5J-L**). Given previous reports that *Gpr18* may also signal through Gαi, we next sought to rule out the involvement of inhibitory G proteins. To this end, we utilized LysM-PTX mice, where myeloid Gi signaling is selectively disrupted (**Figure 5M**). GSIS following NAGly treatment was comparable between LysM-PTX and wild-type controls, reinforcing that *Gpr18* exerts its suppressive effects via Gq, not Gi, signaling in macrophages (**Figure 5N**). There was no difference in body weight observed for any of the cohorts between control and LysM-Gq KO/LysM-PTX(**Figure S7H-I**).

Next, to assess whether the inhibitory effect of GPR18 activation could be attributed to intraislet macrophage populations, we performed static GSIS assays using isolated islets from wild-type and LysM-Gq KO mice. NAGly treatment significantly blunted glucose-induced insulin secretion in wild-type islets, whereas no such suppression was observed in islets derived from LysM-Gq KO animals (**Figure 6A**). We further tested whether depletion of islet macrophages would alter the response. To this end, clodronate and vehicle liposomes were administered to HFD-fed mice, and islets were isolated and subjected to GSIS. As expected, islets from animals treated with clodronate did not exhibit NAGly-induced inhibition of insulin secretion, demonstrating that this suppressive effect is mediated predominantly by resident macrophages (**Figure 6B**). To further validate these findings in a co-culture model, conditioned media from wild-type or Gq KO BMDMs cells were treated with NAGly and incubated with INS-1 β-cells. Only supernatants from wild-type macrophages suppressed insulin release (**Figure 6C**), confirming the requirement of myeloid Gq signaling for GPR18-mediated β-cell inhibition. However, no significant difference was observed for glyburide, arginine and exendin 4 when INS-1 cells were treated with the supernatants (**Figure S7J-L**).

**Figure 6.**
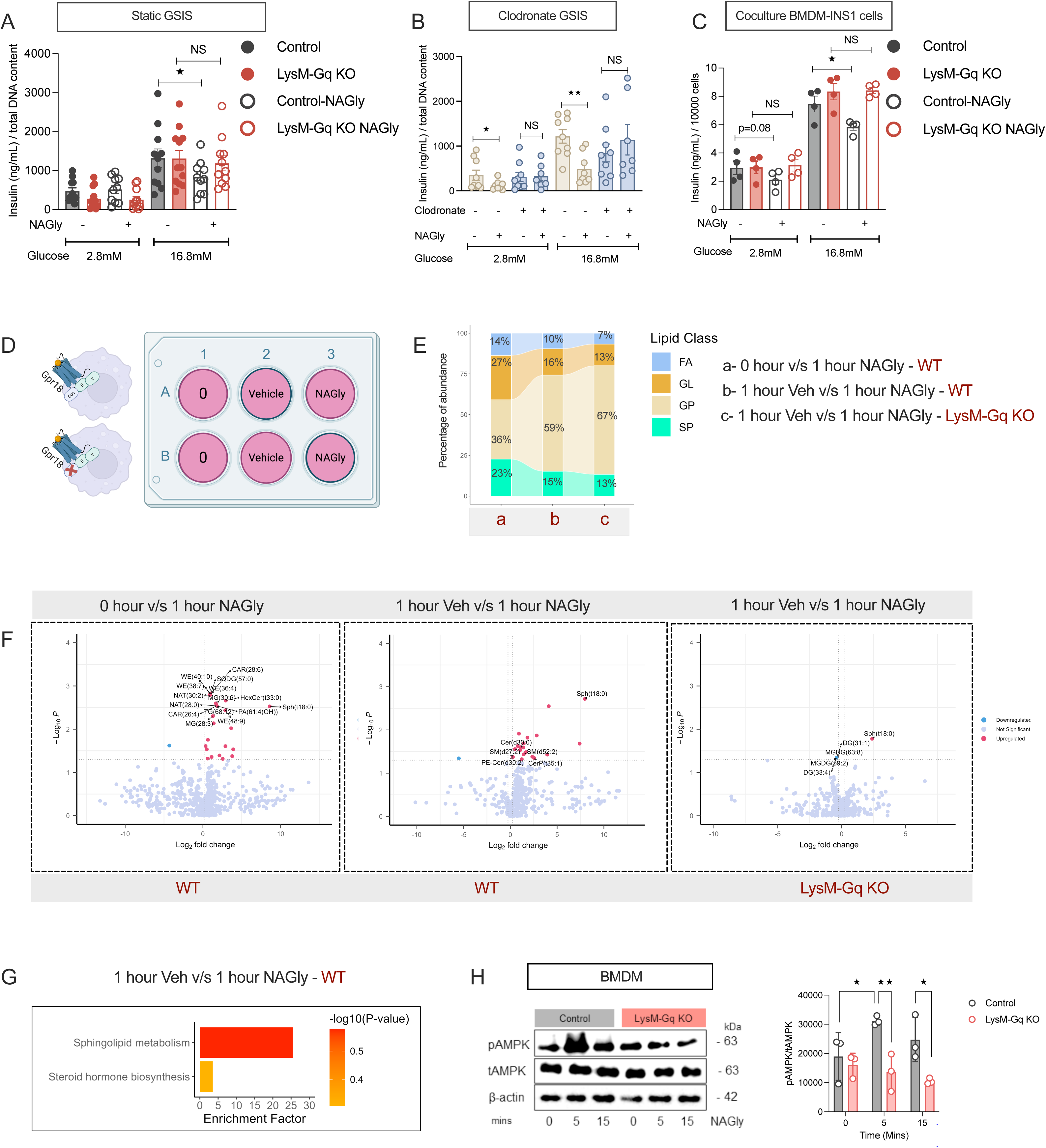
Macrophage GPR18 activates the Gq–AMPK pathway to promote sphingolipid release. (A) Static GSIS from islets of LysM-Gq KO and control mice stimulated with low (2.8 mM) or high (16.8 mM) glucose (n=10-12) in presence or absence of NAGly (10 µM). (B) Static GSIS in clodronate-treated (macrophage-depleted) islets under low and high glucose with or without NAGly (10 µM; n=7-9). (C) Co-culture of BMDMs with INS-1 cells showing insulin secretion at low (2.8mM) and high (16.8mM) glucose in the presence or absence of NAGly (10 µM; n=4). (D) Experimental schematic of WT and LysM-Gq KO macrophages treated with vehicle or NAGly for 1 hour, alongside unstimulated controls (n=). (E) Stacked area plots of lipidomic profiles showing relative abundance of fatty acids (FA), glycerolipids (GL), glycerophospholipids (GP), and sphingolipids (SP) across conditions: (a) WT, 0 hour vs 1 hour NAGly; (b) WT, 1 hour vehicle vs 1 hour NAGly; (c) LysM-Gq KO, 1 hour vehicle vs 1 hour NAGly. Percentages represent the fraction of each lipid class within total lipid species. (F) Volcano plots of differential lipid species (log₂ fold change vs –log₁₀ p-value) after NAGly treatment: WT, 0 hour vs 1 hour(left); WT, vehicle vs 1 hour NAGly (middle); LysM-Gq KO, vehicle vs 1 hour NAGly (right). (G) Pathway enrichment analysis in WT BMDMs (vehicle vs 1 hour NAGly; 10 µM), highlighting sphingolipid metabolism as the top enriched pathway. (H) Immunoblot and quantification of AMPK phosphorylation in control and LysM-Gq KO BMDMs after NAGly treatment (10 µM; 0–15 min). Lipidomic experiments were done in 3 biological replicates and 2 technical replicates. Data are mean ± s.e.m. using one-tailed unpaired t-test (A-C, H); *p < 0.05, **p < 0.01.

### Macrophage GPR18 activates the Gq–AMPK pathway to promote sphingolipid release

To dissect the mechanism by which GPR18 mediates its effects, we first tested whether its activation recapitulates the lipid signaling observed with chemogenetic acute Gq activation. BMDMs from WT mice were treated with the GPR18 agonist NAGly or vehicle, and cell pellet were collected after 60 minutes for lipidomic analysis (**Figure 6D**). PLSDA of NAGly treatment v/s vehicle and WT v/s LysM-Gq KO stimulated with NAGly substantially altered the BMDM lipidome (**Figure S8A-B**). Similar to GqD-expressing macrophages, NAGly stimulation in WT BMDMs led to a marked increase in sphingolipid species (**Figure 6E, , S5B,H**). Strikingly, this lipid release was abolished in macrophages from myeloid-specific Gαq knockout (LysM-Gq KO) mice, confirming that GPR18 requires Gq signaling to drive sphingolipid secretion and suppress insulin secretion (**Figure 6E-F**). Further, pathway analysis confirmed the enrichment of the sphingolipid metabolism pathway in NAGly treated WT cells (**Figure 6G**). Next, to delineate the downstream signaling, we examined AMPK phosphorylation status by immunoblotting. In WT macrophages, NAGly treatment induced a rapid increase in pAMPK levels within 5–15 minutes, whereas this response was completely abolished in mGq-KO cells (**Figure 6H**). These findings indicate that GPR18 utilizes the same Gq–AMPK–lipid axis as chemogenetic GqD activation.

To determine whether this signaling axis is conserved in human immune cells, we took advantage of human islet scRNA seq data and looked for *GPR18* expression and *GPR18* was expressed in immune cells including macrophages (**Figure 7A**). Next, to confirm these observations, we differentiated PBMCs into macrophages and assessed GPR18 expression and *GPR18* levels positively correlated with BMI (**Figure 7B**).While *GPR18* expression trended higher in pro-inflammatory M1 macrophages compared with M0 controls (**Figure 7C**). Functionally, NAGly stimulation of human macrophages increased AMPK phosphorylation, mirroring the response observed in murine cells **(Figure 7D**). To further investigate the lipidomic profile of human PBMCs, we activated GPR18 using NAGly and inhibited downstream signaling through Gαq (UBO) and AMPK (CpdC) **(Figure 7E)**. PLS-DA analysis comparing untreated and NAGly-treated cells, with or without inhibitor treatment, demonstrated that NAGly stimulation significantly altered the PBMC lipidome, whereas these changes were substantially diminished in the presence of UBO or CpdC **(Figure S8C–E)**. Specifically, NAGly treatment led to a marked increase in sphingolipid abundance; however, this effect was abolished when GPR18 activation occurred concurrently with inhibition of Gαq or AMPK **(Figure 7F)**. Consistent with these findings, pathway enrichment analysis revealed that NAGly-induced activation of GPR18 upregulated sphingolipid metabolic pathways, while inhibition of these intermediary signaling nodes prevented this metabolic shift **(Figure 7G–I)**.

**Figure 7.**
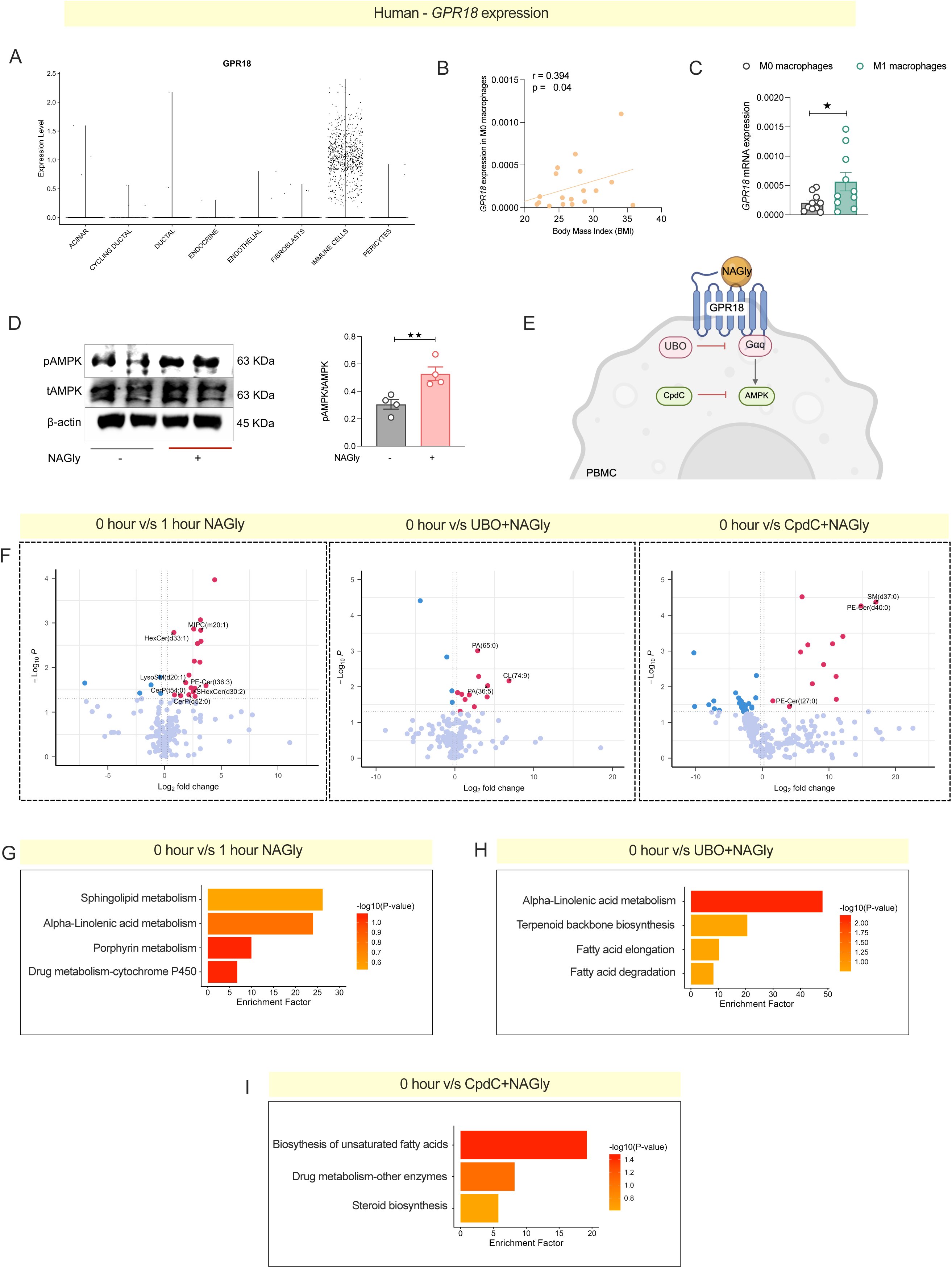
Human PBMC GPR18-Gq–AMPK pathway to promote sphingolipid metabolism. (A) Violin plot of scRNA seq data showing *GPR18* expression across pancreatic islet cell types. (B) Correlation of *GPR18* mRNA expression in M0 macrophages with body mass index (BMI). (C) *GPR18* mRNA expression in PBMC-derived macrophages under M0 and M1 polarization, showing higher levels in M1 macrophages. (D) Western blot analysis of pAMPK activation upon NAGly treatment in PBMCs (n=4). (E) Schematic of Gpr18 activation via NAGly and downstream inhibition of Gaq with UBO and AMPK with CpdC. (F) Volcano plots of differential lipid species (log₂ fold change vs –log₁₀ p-value) after NAGly treatment in PBMC: 0 hour vs 1 hour NAGly(left); 0 hour vs UBO+NAGly (middle); (H) 0 hour vs CpdC+NAGly (right). (G-I) Pathway analysis of differential lipid species (log₂ fold change vs –log₁₀ p-value) after NAGly treatment in PBMC: 0 hour vs 1 hour NAGly(G); 0 hour vs UBO+NAGly (H); (H) 0 hour vs CpdC+NAGly (I).(D-I) Lipidomic experiments were done in 3 biological replicates and 2 technical replicates. Data are mean ± s.e.m. using one-tailed unpaired t-test (A-D); *p < 0.05, **p < 0.01.

Collectively, these data demonstrate that both mouse and human macrophages signal through a conserved GPR18–Gq–AMPK pathway that promotes sphingolipid release and impairs β-cell insulin secretion **(Figure 8)**.

**Figure 8.**
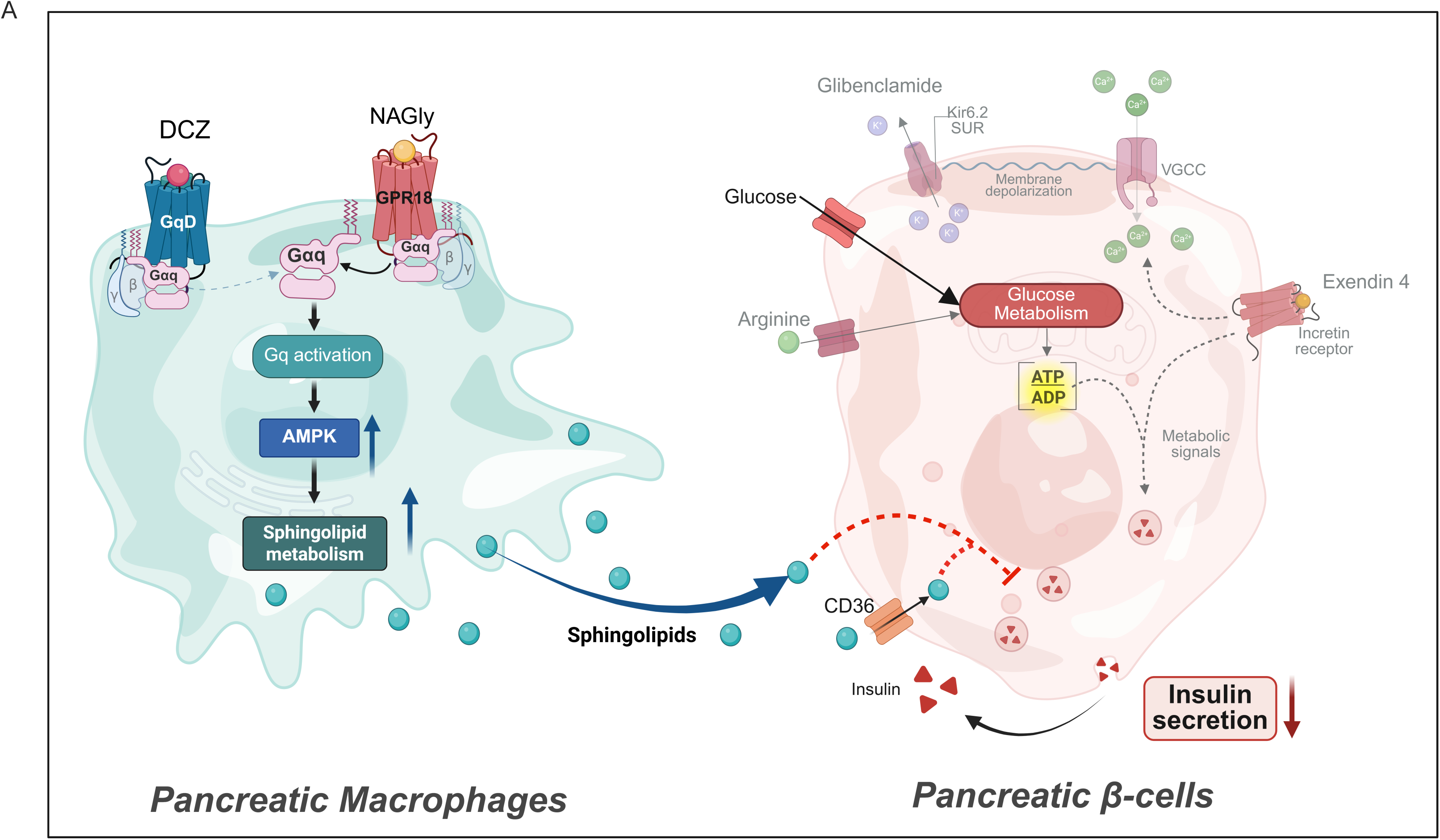
Summary schematic of proposed mechanism. Activation of Gq signaling in macrophages through the GqD/GPR18 pathway, increased phosphorylation of AMPK leading to the stimulation and release of sphingolipids and their derivatives into the extracellular environment. These bioactive sphingolipids act as signaling molecules that enter pancreatic β-cells via the CD36 transporter, where they inhibit insulin secretion through alternative intracellular signaling cascades.

## Discussion

Most tissue-resident macrophages originate during embryogenesis and act as highly specialized immunometabolic sentinels, integrating nutritional, lipid, and metabolic signals to regulate both immune tone and local metabolic programs [6, 29, 30]. Beyond producing cytokines, macrophage generates bioactive metabolites that influence stress adaptation and cellular homeostasis. Although several lipid mediators have been implicated in chronic β-cell dysfunction [31–33], the upstream immune signals that drive their release, particularly under acute conditions, remains poorly understood.

In the current study, we identified macrophage Gq signaling as a dynamic regulator of β-cell function and systemic glucose homeostasis. We demonstrate that acute activation of myeloid Gq signaling rapidly impairs glucose clearance by suppressing β- cell insulin secretion. Mechanistically, Gαq-stimulated macrophages secrete sphingolipids, including ceramides, that disrupt β-cell insulin signaling. Conversely, myeloid deletion of Gαq enhances GSIS and glucose tolerance. Finally, we identify GPR18 as a Gq-coupled receptor in macrophages that mediates this effect, and its agonist *N*-arachidonoyl glycine (NAGly) recapitulates β-cell suppression *in vivo*, but only if macrophage Gq is intact. These findings identify macrophage Gq signaling, specifically the GPR18–Gq–sphingolipid axis, as a key regulator of immune–islet crosstalk in metabolic disease.

Our chemogenetic LysM–GqD model revealed that acute Gq activation in myeloid cells impaired glucose tolerance and GSIS without altering insulin sensitivity in ITTs (**Figure 1**). This divergence from canonical obesity-induced inflammation models suggests macrophage Gq signaling primarily targets the insulin secretion axis rather than peripheral insulin resistance. This selective suppression of β-cell secretion was evident across multiple secretagogues, including glucose, glyburide, and arginine stimulation in Gq-activated mice. Interestingly, while exendin-4 (a GLP-1 agonist) elicited preserved insulin release under a chow diet despite macrophage Gq activation, the same response was blunted in HFD-fed mice, indicating that obesity sensitizes β-cells to this inhibitory pathway (**Figure 2**). Mechanistically, the inhibition occurred upstream of Ca²⁺ entry, since it was bypassed by directly opening L-type Ca²⁺ channels with FPL64176 in co-culture experiments. Thus, the β-cell Ca²⁺ entry and exocytotic machinery remain intact, but a macrophage-derived factor suppresses depolarization-driven responses, which is a hallmark of β-cell lipotoxicity and inflammatory inhibition [34]. Our LysM-GqD model demonstrates that activating a single signaling node in macrophages is sufficient to phenocopy these effects acutely. Conversely, LysM-Gq knockout mice had enhanced glucose clearance and a trend towards improved GSIS without changes in weight or insulin sensitivity (**Figure 4**). Thus, both gain and loss-of-function of macrophage Gq produce reciprocal effects on insulin secretion and glucose tolerance, underscoring a causative role.

Indirect co-cultures of murine macrophages with rat insulinoma cells recapitulated these inhibitory effects, pointing to soluble mediators as the driver. Lipidomics revealed a rapid (<1 h) increase in secreted sphingolipids following Gq activation. This release was AMPK-dependent, as compound C blocked sphingolipid production and preserved β-cell insulin secretion. Functionally, these sphingolipids traversed β-cell membranes and inhibited Akt phosphorylation, a known molecular mechanism for ceramide-induced insulin resistance [35, 36]. Notably, while previous work has implicated adipose, liver, and muscle ceramide pools in systemic insulin resistance [37, 38], our data position islet macrophages as a rapid, local source of sphingolipids capable of suppressing β-cell function independent of chronic inflammation. Ceramides impair insulin signalling by activating PP2A, which dephosphorylates Akt, and PKCζ-mediated inhibition of Akt membrane translocation [36]. In humans, CERS6, the primary enzyme responsible for synthesizing C14–C16 ceramides, correlates with BMI, hyperglycemia, and decreased insulin sensitivity. Its inhibition leads to improved metabolic health [36]. Notably, while ceramide accumulation is typically considered a chronic process during overnutrition, our findings showed for the first time that acute Gq signaling activation releases ceramides and influences the β-cell function. Previous studies have shown that ceramides can signal through FPR2 in adipocytes [39], raising the possibility that a similar mechanism might contribute to β-cell inhibition. However, scRNA-seq analysis of islet cell populations did not detect FPR2 expression in β cells (data not shown), suggesting that alternative pathways mediate the effects of ceramides in the islet microenvironment.

We further identified GPR18 as a physiological receptor mediating this pathway. Pharmacological activation of GPR18 with the agonist *N*-arachidonoyl glycine reproduced the effects of Gq-DREADD stimulation. These responses were absent in LysM–Gq knockout mice, demonstrating a requirement for myeloid Gq signaling. These findings are consistent with broader evidence that Gq-linked GPCR signaling in macrophages can rapidly remodel the secretome through Ca²⁺- and lipid-dependent mechanisms [40]. Our findings extend prior work on islet macrophage GPCRs, where FFAR4, another Gq-coupled receptor, promotes GSIS via IL-6 secretion under physiological conditions [15], while GPR92 deficiency impairs insulin secretion and enhances islet inflammation through cytokine-mediated mechanisms [14]. In contrast, Gq/GPR18 signaling suppresses GSIS via lipid mediators, revealing a new immune–islet communication axis. Our identification of GPR18 as a Gq-coupled receptor that suppresses β-cell insulin secretion adds a new layer to its known roles. In cardiovascular and liver systems, activation of myeloid GPR18 by the specialized pro-resolving mediator resolvin D2 (RvD2) promotes efferocytosis, increases reparative Arg1⁺ macrophages, and limits plaque necrosis and fibrosis[41, 42]. These effects arise alongside an anti-inflammatory lipid profile enriched in specialized pro-resolving mediators (SPMs). In contrast, our work shows that acute Gq-biased activation of GPR18 in islet macrophages triggers rapid AMPK-dependent release of sphingolipids, including ceramides. Whereas SPMs promote repair and metabolic balance, ceramides impair insulin signalling and blunt insulin secretion. This highlights a ligand- and context-dependent switch in GPR18 outputs.

Our study uncovers a macrophage-to-β-cell communication axis mediated by Gq signaling and lipid secretion, adding a new layer to the concept of “metaflammation” in diabetes. Whereas, previous work has emphasized on cytokines (IL-1β, TNFα, IL-6) and cell–cell contact as macrophage-derived inhibitors of insulin secretion. This study added lipid mediators, specifically ceramides, as paracrine messengers linking macrophage Gq activation to β-cell dysfunction. In obesity and T2D, increased availability of fatty acids and endocannabinoid metabolites (like NAGly) could chronically engage this pathway, contributing to β-cell failure. Conversely, dampening macrophage Gq activity by targeting the GPR18–Gq axis could preserve β-cell insulin secretion and improve glucose homeostasis. Future studies should also examine how chronic Gq activation reshapes macrophage phenotype and whether this signaling intersects with other inflammatory pathways, such as the NLRP3 inflammasome or TLR signaling, during obesity.

Finally, the translational relevance is underscored by our observation that GNAQ expression in human PBMCs correlates with BMI, suggesting that this mechanism may also operate in humans. A previous study reported that arachidonic acid containing ether lipids are enriched in obesity, likely reflecting alterations in lipid metabolic pathways [43]. Future work will need to assess macrophage-derived sphingolipids and GPR18 signaling in human islets and to explore therapeutic approaches targeting this pathway. By establishing macrophage Gq signaling as a central regulator of β-cell function, our study provides a new conceptual framework for understanding immune–islet crosstalk in obesity and diabetes.

### Limitation of the study

While we found sphingolipids dysregulation in macrophages regulates insulin secretion from β-cell, all mechanistic studies were conducted in mouse models, which may not fully reflect the situations in human islets. Further human studies, including validating the sphingolipids signaling in islet samples from human, performing lipidomics from islet isolated macrophages are needed to fully validate our proposed mechanism.

## Methods

### Drugs, reagents, commercial kits, and antibodies

The sources of all drugs, reagents, commercial kits, antibodies, and mouse strains are detailed in Resource Table 1.

### Generation and maintenance of mutant mice

Mutant mice expressing a Gq-coupled DREADD (hM3Dq) [17]selectively in myeloid cells (LysM-GqD mice) were generated by crossing LysM-Cre mice (The Jackson Laboratory, stock # 018956) with LSL-hM3Dq mice (The Jackson Laboratory, stock # 026220) . Mice carrying one copy of the LSL-hM3Dq gene but not the LysM-Cre transgene served as control animals. To generate myeloid-cell-specific Gaq/11 KO mice (LysM-GqKO), Gna11^-/-^ Gnaq^fl/fl^ mice [44, 45] were crossed with LysM-Cre mice, resulting in the experimental cohort of LysM-Cre-Gna11^-/-^Gnaq^fl/fl^ mice. Cre-negative Gna11 ^-/-^ Gnaq^fl/fl^ mice were utilized as control mice in all knockout experiments. For selective disruption of Gαi signaling in myeloid cells, ROSA26-PTX^fl/fl^ (RRID:MMRRC_030678-UCD) mice [46, 47], carrying a floxed-stop cassette upstream of the gene encoding the catalytic subunit of pertussis toxin (S1-PTX), were crossed with *LysM-Cre* mice. The resulting *LysM-Cre ROSA26-PTX^flox/+^* offspring are referred to as *LysM-PTX* mice, with Cre-negative *ROSA26-PTX^flox/+^* littermates serving as controls. All mouse strains were maintained on a C57BL/6NJ background.

All mice were housed at 23°C within a pathogen–free facility that featured a 12- hour light and 12-hour dark cycle, with unrestricted access to regular chow (RC diet with 12.6 kcal% Fat Safe Diet D131) and water. In a subset of experiments, mice (age: 6-8 weeks) were maintained on a high-fat diet (HFD) (F3282, 60% kcal fat, energy density 5.5 kcal/g, Bioserv) for a minimum duration of 8 weeks prior to conducting any metabolic studies. Unless otherwise specified, adult male littermates were utilized for all experiments. The wild-type mice used for some of the studies were acquired from Jackson Laboratory (C57BL/6NJ mice). Body weight and chow intake were assessed on a weekly basis. The research methods described in this manuscript adhere to all pertinent ethical regulations. All animal experiments received approval from the Institutional Animal Ethics Committee (IAEC) at the Indian Institute of Technology Kanpur, India.

All animals used in this study were littermate-controled male mice on a pure C57BL/6NJ background. The age and number of the mice used for each experiment are indicated in the figure legends.

### In vivo metabolic test

#### Deschloroclozapine (DCZ) treatment of LyzM-GqD mice

LyzM-GqD and control littermate mice, maintained on either an RC or an HFD, were subjected to a six-hour fast (from 8:00 am to 2:00 pm), followed by the intraperitoneal administration of 30 µg/kg body weight of DCZ. Blood glucose levels were measured from the tail vein with a glucometer (Contour Next; Ascensia Diabetic Care) at 0, 15, 30, 60, 90, and 120 minutes post-injection. Similarly, to measure plasma insulin and non-esterified fatty acids (NEFA), experimental and control mice received 30 µg/kg body weight of DCZ after 6 hours of fasting. Plasma samples were collected at 0, 15, 30, and 60 minutes post-injection, and the plasma analytes were measured following the manufacturer’s protocol.

#### Glucose, insulin, and pyruvate tolerance test

Glucose tolerance test (GTT), insulin tolerance test (ITT), and pyruvate tolerance test (PTT) were conducted on mice aged ≥8 weeks that were either maintained on a regular chow (RC) diet or fed a high-fat diet (HFD) for at least 8 weeks. For all metabolic tests, DCZ was administered intraperitoneally (i.p.) at 30 µg/kg body weight 45 minutes prior to glucose, insulin, or pyruvate injection in both experimental (LysM-GqD) and control cohorts. For RC-fed mice, an additional condition involved a lower dose of DCZ (10 µg/kg) administered 45 minutes before glucose injection (2 g/kg body weight). GTTs were conducted following an overnight fast (8:00 PM–9:00 AM) where mice received either 2 g/kg glucose (RC condition) or 1 g/kg (HFD condition) glucose either i.p. or by oral gavage. For ITT and PTT, the mice were fasted for six hours (from 8:00 AM to 2:00 PM) and insulin (Humulin, Eli Lilly) or sodium pyruvate was administered intraperitonially. For the ITT, insulin dosage of 0.75 U/kg was administered to the RC mice, while a dosage of 1 U/kg was administered to the HFD mice. In the case of PTT, 2 g/kg of sodium pyruvate was utilized for the RC mice, and 1 g/kg was administered to the HFD-fed mice. Then the blood glucose levels were measured from the tail vein using a glucometer (Contour Next; Ascensia Diabetes Care) at the following time points: –45, 0, 15, 30, 60, 90, and 120 minutes. For statistical analyses, the area under the curve (AUC) was determined after subtracting baseline glucose levels. Initial comparisons were made using Student’s t-test; follow-up comparisons at individual time points were performed with Student’s t-test with Bonferroni correction.

#### Glucose/ Glyburide/ Exendin 4/ Arginine/Oral glucose stimulated insulin secretion

All experiments were conducted on mice fasted for six hours (8:00 AM–2:00 PM). DCZ was administered intraperitoneally at a dose of 30 µg/kg body weight 45 minutes prior to the administration (i.p. or oral gavage) of glucose, glyburide, exendin-4, or arginine. For glucose-stimulated insulin secretion (GSIS) and oral GSIS (O-GSIS) assays, RC-fed mice received 2 g/kg glucose and HFD-fed mice received 1 g/kg glucose [22]. Glyburide was administered i.p. at 5 mg/kg and 10 mg/kg for RC and HFD groups, respectively [48, 49]. Exendin-4 was given at 12 nmol/kg for RC-fed mice and 4 nmol/kg for HFD-fed mice. Arginine was administered at 1 g/kg body weight for both RC and HFD groups [50]. Blood samples were collected from the tail vein into heparin-coated tubes for hormone and metabolite analysis. Plasma was separated by centrifugation at 10,000 × g for 10 minutes at 4 °C and stored at −80 °C until analysis. Plasma insulin levels were measured using commercial ELISA kits (Crystal Chem, and ALPCO, USA).

### Bone marrow-derived macrophages (BMDM) culture

BMDMs were derived from male and female mice aged between 8 and 12 weeks and were maintained on either RC or HFD, per established protocols [4]. In brief, Bone marrow was flushed from femurs and tibias using ice-cold DMEM medium, passed through a 100 µm cell strainer, and centrifuged at 2,000 RPM for 5 minutes at 4 °C. Cells were cultured in complete medium supplemented with 20 ng/mL macrophage colony-stimulating factor (M-CSF) for four days, after which the medium was replaced with fresh M-CSF-supplemented medium. Cells were used for downstream assays from day 6 onward.

### Immunostaining

Pancreata were harvested following 0.9% saline perfusion and fixed in 4% paraformaldehyde (PFA) at 4°C for 12 hours. Fixed tissues were dehydrated through a graded ethanol series (25%, 50%, 70%, 90%, and 100%), cleared in xylene, and embedded in paraffin. Serial sections (10 µm thick) were cut using a microtome for histological evaluation.

For immunofluorescence staining, sections were deparaffinized, rehydrated, and subjected to antigen retrieval as per standard protocols [51]. To assess colocalization of the GqD receptor with macrophages, sections were incubated with primary antibodies against the HA tag (to detect GqD) and F4/80 (a macrophage marker), followed by Alexa Fluor 488-conjugated anti-rabbit secondary antibody for HA and Alexa Fluor 594- conjugated anti-Rat antibody for F4/80. Slides were mounted with antifade medium containing DAPI and imaged using a fluorescence microscope (Nikon ECLIPSE Ti2-E inverted microscope).

### Islet isolations and culture

Pancreata were harvested from mice following euthanasia as per institutional ethical guidelines. The abdominal cavity was exposed via midline laparotomy, and the pancreas was carefully located and isolated to avoid damage to surrounding tissues. For islet isolation, the pancreas was perfused through the common bile duct with a collagenase solution (3.5 mg/mL in Hanks’ Balanced Salt Solution (HBSS) without calcium and magnesium) using a 30G needle, with the hepatopancreatic ampulla clamped to ensure targeted perfusion. The pancreas was then carefully dissected and incubated at 37°C for 10-15 min for enzymatic digestion. Following digestion, the tissue was neutralized using wash buffer (HBSS ( pH 7.4) supplemented with 0.02% BSA, 1× Pen/Strep, and 5 mM HEPES and dispersed by gentle trituration. The suspension was subjected to density gradient separation by layering over Histopaque-10771 (Sigma) and centrifuged at 450 g for 20 minutes at 4°C. Purified islets were manually picked under a stereo microscope and cultured overnight in islet culture medium (RPMI-1640 supplemented with 10% fetal bovine serum (FBS), 1% penicillin-streptomycin, and 5.5 mM glucose) at 37°C in a humidified 5% CO₂ atmosphere.

### Static glucose stimulated insulin secretion

Post islet isolation, islets were incubated in islet culture medium (RPMI-1640 supplemented with 10% fetal bovine serum (FBS), 1% penicillin-streptomycin, and 5.5 mM glucose) at 37°C in a humidified 5% CO₂ atmosphere for 6 hours. Post six hours of incubation, the islets were incubated in KRBH (NaCl-128.8mM, KCl-4.8mM, KH2PO4- 1.2mM, MgSO_4_- 1.2mM, CaCl_2_ – 2.5mM, NaHCO_3_ – 5mM, HEPES – 10mM, BSA – 0.1%) buffer without glucose for 30 min and then stimulated with DCZ (30nM)/NAGly (10 µM) for 45 min in 2.8mM glucose. After 45 mins, supernatant was collected and stimulated in 16.8mM glucose for 30 mins[52]. Supernatant was collected and ELISA was performed using ALPCO STELLUX Chemiluminescence kit. Data was normalized with the total DNA content of the islets.

### BMDM and INS-1 cells culture and coculture

INS-1 cells (6 × 10⁴/well) and BMDM (1 × 10⁴/well) were seeded in 96-well plates. On day 4, macrophages were starved for 60 min in modified Krebs–Ringer Bicarbonate HEPES (KRBH) buffer was prepared containing 2.54 mM CaCl₂, 1.19 mM MgSO₄, 1.199 mM KH₂PO₄, 4.64 mM KCl, 25 mM NaHCO₃, and 10 mM HEPES, and INS-1 cells were starved for 15 min in basal RPMI with 1 mM glucose. BMDMs were then treated with 30 nM DCZ for 45 min. Following treatment, 75 µL of macrophage supernatant was transferred to INS-1 cells for 15 min. Cells were subsequently stimulated for 30 min with test compounds in KRBH buffer: low glucose (2.8 mM), high glucose (16.7 mM), glyburide (100 nM), arginine (30 mM), FPL (0.1 µM), or Exendin-4 (10nM). After treatment, supernatants were collected without dilution and assayed by ELISA using ALPCO STELLUX Chemiluminescence kit.

To validate that sphingolipid uptake was facilitated by CD36, INS-1 cells were incubated (30 min at 37°C) with 0.5 mmol/l of the irreversible CD36 inhibitor, sulfosuccinimidyl-oleate (SSO), and then washed twice (KRBH buffer with 0.2% BSA) and assayed for GSIS.

### qRT-PCR analysis of gene expression

Total RNA was extracted from frozen tissues or cultured cells using the RNA extraction kit (Promega) in accordance with the manufacturer’s protocol. Complementary DNA (cDNA) was synthesized using the cDNA synthesis kit (Takara). Quantitative PCR (qPCR) was performed using the SYBR Green method (Takara). Gene expression data were normalized to 18S rRNA expression using the ΔΔCt method. The primer details are provided in the Resource Table 1.

### Western blot analysis

Protein lysates were prepared by homogenizing the islets and cells in RIPA lysis buffer (25 mM Tris-HCl pH 7.6, 150 mM NaCl, 5 mM EDTA, 1% NP-40 or 1% TritonX-100, 1% sodium deoxycholate, 0.1% SDS) supplemented with protease and phosphatase inhibitors (Roche 4906845001). The lysate was collected through centrifugation at 15,000 RPM at 4°C , and protein concentrations were determined using the Pierce BCA Protein Assay Kit (Thermo Fisher Scientific). Proteins were resolved in the Tris-glycine buffer and transferred to PVDF membranes using the BioRad semi-dry transfer system (BioRad). The blots were then blocked with 5% BSA for 1 hour at room temperature and then incubated with primary antibodies (1:1000) diluted in 5% BSA blocking buffer overnight at 4°C. The membranes were then washed with 1X TBST and incubated with HRP-conjugated secondary antibody (1:10,000), followed by washing with 1X TBST. The membranes were then developed and imaged with the Azure imaging system using Clarity Max ECL Western Blotting Substrates (BioRad). Finally, images were analyzed and quantified using ImageJ (Fiji).

### Indirect calorimetry and energy expenditure measurements

Energy expenditure measurements were carried out with mice at 23°C using an Oxymax/CLAMS monitoring system (Columbus Instruments). Mice consuming RC and HFD were acclimatized in metabolic chambers for 24 hours and then injected with two injections of DCZ (30 µg/kg, ip) at 8:00 a.m. and 8:00 p.m. the next day. Food intake, water intake, O_2_ consumption, CO_2_ production, and ambulatory activity were measured at 5-min intervals. After a single DCZ injection, mice were monitored for one day.

### Clodronate liposomes and delivery

Clodronate liposomes were prepared as previously described [53]. In brief, phosphatidylcholine (86 mg) and cholesterol (8 mg) were dissolved in chloroform, and the solvent was evaporated under reduced pressure to form a lipid film. The film was hydrated with 10 mL of 0.6 M clodronate (2.5 g in distilled water) and gently rotated under nitrogen at room temperature for 2 hours. The suspension was sonicated (3 min), incubated under nitrogen for an additional 2 hours, and centrifuged at 10,000 × g to remove unencapsulated drug. Liposomes were washed 2–3 times in sterile PBS (25,000 × g, 30 min) and resuspended in 4 mL sterile PBS. Mice were injected intravenously with clodronate liposomes at a dose of 10 µL/g body weight for macrophage depletion.

### Untargeted lipidomics

#### Cell pellet

Bone marrow–derived macrophages (BMDMs) were plated at 0.5 × 10⁶ cells per well. Cells were serum-starved for 1 hour prior to treatment with 30 nM DCZ for either 60 or 120 min. Following treatment, cells were washed twice with 1× PBS and scraped into PBS. Lipids were extracted by adding 2 mL chloroform and 1 mL methanol, followed by vortexing and centrifugation at 2800 g for 5 min. The organic (lower) phase was collected and supplemented with 50 µL formic acid. Samples were then dried under a stream of nitrogen and stored for downstream analysis.

### Cell supernatant

For lipid extraction, 500 μl of cell culture supernatant was mixed with 1.5 mL methanol in a glass tube with a Teflon-lined cap and vortexed. Subsequently, 5 mL MTBE was added, and the mixture was incubated for 1 hour at 4^0^C with shaking. Phase separation was induced by adding 1.25 mL MS-grade water, followed by 10 min incubation at room temperature and centrifugation at 1,000 g for 10 min. The upper organic phase was collected, while the lower phase was re-extracted with 2 mL of a solvent mixture mimicking the expected organic composition [MTBE/methanol/water (10:3:2.5, v/v/v), upper phase]. The combined organic extracts were dried under N_2_ stream.

### LC-MS/MS

To analyze the lipid profiles of the samples, an untargeted lipidomics approach was employed. Lipid extracts were analyzed using an Agilent 1290 Infinity II UHPLC system coupled to an Agilent 6546 quadrupole time-of-flight (QTOF) mass spectrometer. Samples were prepared in methanol:isopropanol (1:1, v/v) and maintained at 4 °C in the autosampler prior to injection. A 5 μL aliquot of each sample was injected into the LC– MS/MS system. Chromatographic separation was performed on an Agilent ZORBAX Eclipse Plus C18 column (2.1 × 50 mm, 1.8 μm) maintained at 60 °C.

### Data analysis

Data Conversion and Preprocessing: Raw files (.d) were converted to mzXML using MSConvert (ProteoWizard, v3.0.25163-1e83dfc) with centroid peak picking (MS1 and MS2). Data were preprocessed in MZmine (v4.7.8): centroid mass detection (noise level = 6.0E2), chromatogram building (min consecutive scans = 5, min absolute height = 5.0E4, m/z tolerance = 0.005 m/z or 10 ppm) and Savitzky–Golay smoothing. Isotopes were grouped (13C filter; m/z tolerance = 0.005 m/z or 10 ppm; RT tolerance = 0.1 min), aligned (Join Aligner; m/z tolerance = 0.005 m/z or 10 ppm; RT tolerance = 0.1 min), and gap-filled (Feature finder; Intensity tolerance = 1%, m/z tolerance = 0.005 m/z or 10 ppm, RT tolerance = 0.1 min).

Feature Annotation: Unified peak lists (CSV, MZmine 2 format) were generated for each dataset. LipidFinder [54] was used in positive ion mode against COMP_DB with all positive ion adducts. Mass tolerance was 0.001 Da for LysM-GqD and 0.005 Da for Gpr18/supernatant datasets. Pathway enrichment analysis was performed using functional analysis [LC-MS] in Metaboanalyst 6.0.

Quality Control and Normalization: Sample normalization (Normalization by sum), Data transformation (log₁₀) and Scaling (Pareto scaling), along with missing value imputation (1/5 of the min. positive value), were performed using Metaboanalyst 6.0 [55].

Statistical Analysis: Replicates were averaged per biological sample. All the statistical analysis including differential features (volcano plot), multivariate analysis (PCA, PLS-DA) was performed using MetaboAnalyst 6.0.

Metaboanalyst statistical analysis Volcano plot thresholds: -0.25 ≤ |log₂FC| ≥ 0.25, P value < 0.05. WGCNA was applied for lipid–lipid network analysis. Heatmaps for differential features were generated in pheatmap; plots in ggplot2/webR/ EnhancedVolcano (R v4.4.1).

### Integration of publicly available scRNA seq datasets for islet cells

To characterize Gpr18 expression in immune cells, we analyzed four publicly available scRNA sequencing datasets: GSE203376, GSE211799 (the integrated MIA), and GSE162512. GSE203376, GSE162512, and our previous dataset encompass RC and HFD dietary intervention, including 1, 4, 8, 16, and 24 weeks of HFD. From GSE211799, we extracted islet cells from db/db models. Within the db/db condition, we grouped samples as control_db/db (chow_WT) and db/db (Sham_Lepr-/-). Overall, the atlas contained 1,96,808 cells and 34220 features. We preprocessed the raw counts, performed cell type annotation using key marker genes. To integrate these datasets, we used scDREAMER’s unsupervised batch integration method [56], where CONDITION (Sham, db/db, HFD, RC) was treated as batches. The resulting latent embeddings were used for clustering the cells using Seurat’s "FindNeighbors" and "FindClusters" functions. We projected the scDREAMER embeddings into two dimensions using the UMAP algorithm to visualize the cells. The resulting clusters were visualized in UMAP space using the "DimPlot" function, grouped by CONDITION and immune cell subpopulation. We examined the expression of key marker genes to assess the molecular signatures of different cell subtypes. Violin plots were generated using the “Vlnplot” function for annotating cell types, and stacked bar plots were plotted from the ggplot2 package. Differential gene expression was performed for the immune cells to show the expression profile of the *Gpr18* gene.

The expression of GPR18 in human pancreatic immune cells was validated using the publicly available dataset GSE229413, considering only the healthy individuals.

### IP1 assay

Receptor-mediated intracellular changes in IP1 second messenger were measured using IP-One Gq kit (Cisbio) using previously established protocol [57]. Briefly, BMDM cells were differentiated in 12-well plates as described before. Cells were serum starved for 1 h and treated with NAGly (10 µM) in stimulation buffer (pH 7.4) containing 50 mM LiCl. After 45 min of stimulation, cells were lysed using 200 µL of lysis buffer (HEPES buffer 1×, 1.5% Triton, O.2% BSA). 14 µL of cell lysate was added to the 384-well low volume white microplates (NEST). Working solutions of IP1-d2 followed by Anti-IP1-Cryptate (3 µL) were added to the samples. Plate was sealed and incubated for 1 hour at room temperature. Fluorescence was measured at 620 nm and 665 nm with the BMG plate reader.

### PBMC culture

Human peripheral blood samples were obtained from healthy donors as per the Institutional Ethics Committee guidelines of the Health Centre, IIT Kanpur. Approximately 20 mL of venous blood was collected into heparinized vacutainers (BD Vacutainer, Sodium Heparin, 158 USP units) and diluted 1:1 with phosphate-buffered saline (PBS). Peripheral blood mononuclear cells (PBMCs) were isolated by density gradient centrifugation using Histopaque PLUS (Sigma-Aldrich) at 1,200 × g for 10 min at room temperature with no brake. The PBMC layer was carefully aspirated, washed twice with PBS (300 × g for 10 min), and subsequently seeded in RPMI-1640 medium (Gibco) for downstream applications, including Western blotting and lipidomic analyses.

### Treatment of mice with a selective Gpr18 receptor agonist

All experiments were conducted on mice fasted for six hours (8:00 AM–2:00 PM). NAGly was administered intraperitoneally at a dose of 36.15 µg/kg 45 minutes prior to the administration (i.p. or oral gavage) of glucose, glyburide, exendin-4, or arginine. For glucose-stimulated insulin secretion (GSIS) and oral GSIS (O-GSIS) assays, RC-fed mice received 2 g/kg glucose and HFD-fed mice received 1 g/kg glucose. Glyburide was administered i.p. at 5 mg/kg and 10 mg/kg for RC and HFD groups, respectively. Exendin- 4 was given at 12 nmol/kg for RC-fed mice and 4 nmol/kg for HFD-fed mice. Arginine was administered at 1 g/kg body weight for both RC and HFD groups. Blood samples were collected from the tail vein into heparin-coated tubes for hormone and metabolite analysis. Plasma was separated by centrifugation at 10,000 × g for 10 minutes at 4 °C and stored at −80 °C until analysis. Plasma insulin levels were measured using commercial ELISA kits (Crystal Chem, and ALPCO, USA).

### Statistics

In all figures, data are presented as means ± SEM, and error bars denote SEM, unless otherwise noted in the legends. Statistical comparisons were performed using unpaired two-tailed Student’s t tests and one-way or two-way analysis of variance (ANOVA), followed by Bonferroni’s or Sidak’s post hoc test for multiple comparisons, as appropriate (GraphPad Prism). A P value of <0.05 was considered significant.

## Acknowledgements

We sincerely thank Prof. Dr. Stefan Offermanns (Max Planck Institute for Heart and Lung Research, Germany) for sharing the Gna11^-/-^ Gnaq^fl/fl^ mice. We are grateful to Dr. Prosenjit Mondal (IIT-Mandi) for generously providing INS-1 cells. The mouse strain B6;129P2-Gt(ROSA)26Sortm1(ptxA)Cgh/Mmucd, (RRID:MMRRC_030678-UCD) was obtained from the Mutant Mouse Resource and Research Center (MMRRC) at University of California at Davis, an NIH-funded strain repository, and was donated to the MMRRC by Shaun Coughlin, M.D., Ph.D., University of California, San Francisco. We also acknowledge the members of the MMCSL laboratory for their valuable discussions and critical input throughout the study. The current study is primarily supported by the Department of Science and Technology - Science and Engineering Research Board (DST-SERB) [CRG/2021/004502] and DBT/Wellcome Trust India Alliance (IA/I/21/1/505613). Research in the SPP laboratory is supported by the Department of Biotechnology (DBT) [B1/PR44526/MED/30/2376/2021], the Indian Council of Medical Research (ICMR) [5/4/8-18/Obs/SPP/2022-NCD-II], and an Initiation Grant from IIT Kanpur. SS acknowledges support from the DBT Junior Research Fellowship (DBTHRDPMU/JRF/BET/-20/1/2020/AL/352). AK acknowledges support from Ministry of human resource development fellowship by Indian Institute of Technology (IITK/21118264). We acknowledge Ms. Raashidha Farhath for helping us with *in vivo* studies. Authors utilized digital tools, including Grammarly and AI-assisted platforms, for the refinement of grammar and sentence construction.

## Author Contribution

Experimental design was carried out by S.S., A.K., L.F.B., D.A., and S.P.P. All *in vivo* studies were performed by S.S., A.K., S.P., H.Y., S.B., S.K., and K.K.P. Lipidomics experiments were conducted by S.S., and S.D., with data analysis performed by M.S., A.G., S.S., A.K., D.A., and S.P.P. *Ex vivo* islet studies were performed by A.K. and S.S. Liposomes were designed and prepared by S.K.M., T.S., and A.K. Immunostaining was performed by A.K. and S.T. Plasma analyses were conducted by S.S., A.K., R.P., and S.A. Single-cell RNA-seq data analysis was carried out by M.S., S.S., and H.Z. The project was conceptualized by S.P.P. The manuscript was written by S.S., A.K., and S.P.P., with input from all authors. All authors reviewed and approved the final version of the manuscript.

## Supplementary Figures

**Supp Figure 1.**
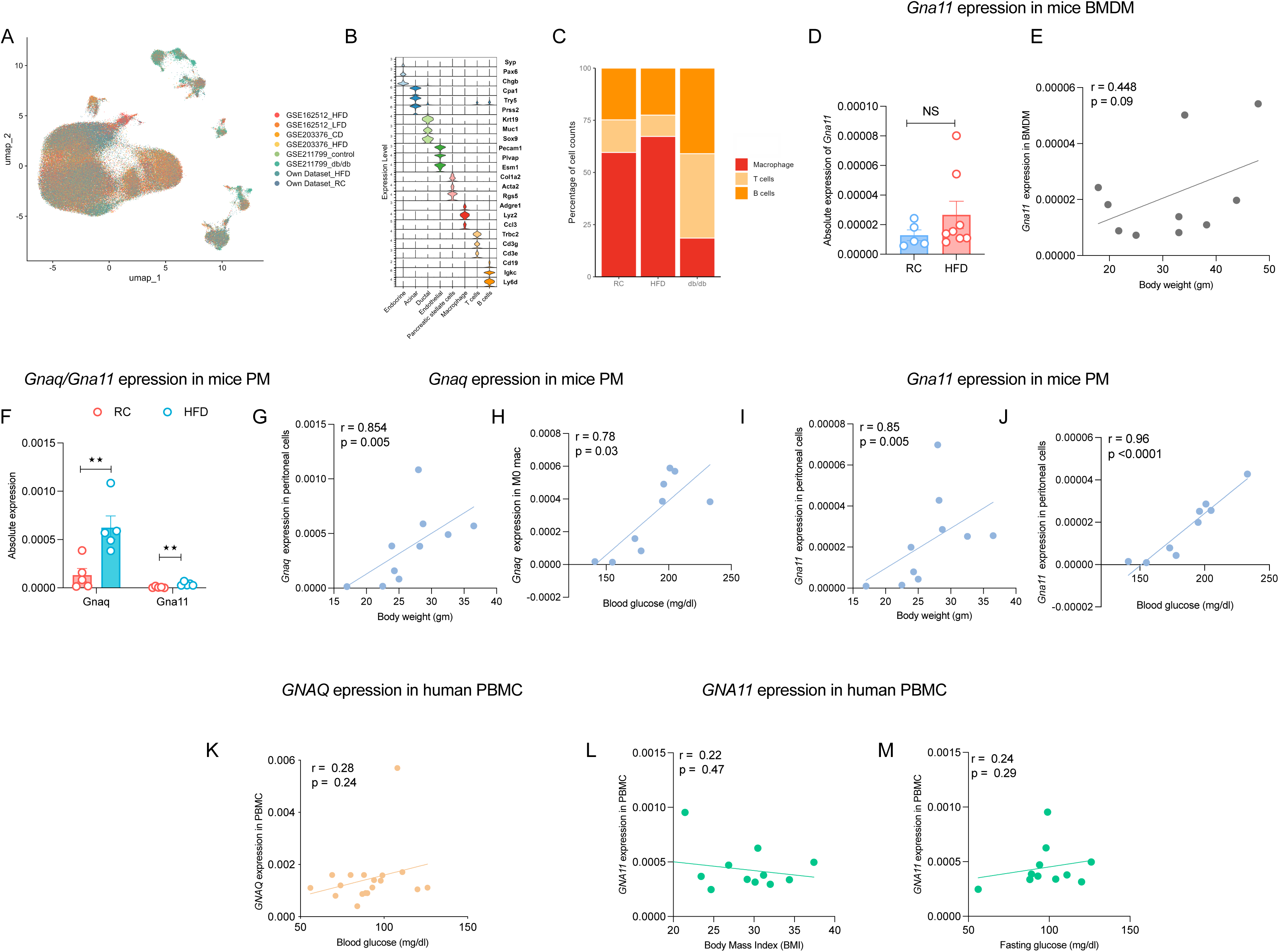
Gq/11 expression in mouse macrophages and human PBMCs. (A) Integrated UMAP of scRNA seq from murine islets from published datasets from RC, HFD and Obese (B) Violin plots of marker gene expression from scRNA seq datasets of islet cell types in mice fed regular chow (RC), high-fat diet (HFD), or db/db background. (C) Major immune cell subsets (macrophages, T cells, B cells) across RC, HFD, and db/db mice, represented as percentage of population. (D) *Gna11* expression in bone marrow–derived macrophages (BMDMs) from RC and HFD mice (n=5-8). (E) Correlation of *Gna11* expression in peritoneal macrophages (PMs) with body weight (n=10). (F) Expression of *Gnaq* and *Gna11* in PMs from RC versus HFD mice (n=5). (G-H) Correlation of *Gnaq* expression in PMs with (G) body weight and (H) blood glucose (n=9-10). (I-J) Correlation of *Gna11* expression in PMs with (I) body weight and (J) blood glucose (n=9-10). (L-N) Human PBMC analysis: *GNAQ* expression correlated with (L) blood glucose, and *GNA11* expression with (M) body mass index (BMI) and (N) fasting glucose (n=12-18). Data represented as mean ± s.e.m.; Pearson’s correlation shown for E, G–J, K–N. *p < 0.01, p < 0.05, NS = not significant.

**Supp Figure 2.**
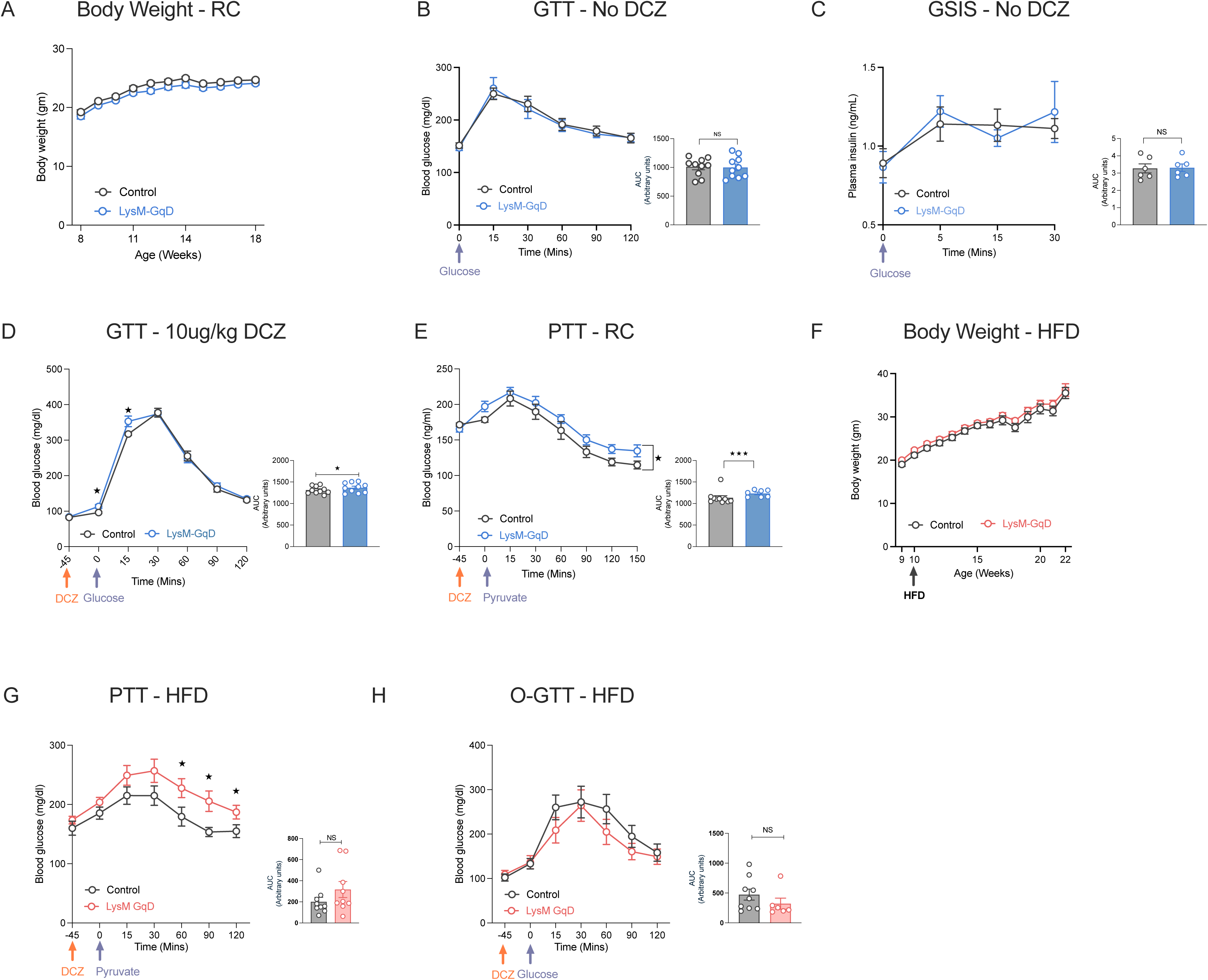
Glucose homeostasis and hormone regulation in LysM-GqD and control mice under regular chow (RC) and high-fat diet (HFD) condition. (A) Body weight of LysM-GqD and control mice maintained on RC (n=7-11). (B) Intraperitoneal glucose tolerance test (GTT) (n=10) and (C) Glucose-stimulated insulin secretion (GSIS) in RC-fed LysM-GqD and control mice without DCZ stimulation (n=6). (D) GTT in RC-fed LysM-GqD and control mice after acute DCZ (10 µg/kg) stimulation (n=10). (E) Pyruvate tolerance test (PTT) in RC-fed LysM-GqD and control mice with DCZ (30 µg/kg) stimulation (n=7-11). (F) Body weight progression of LysM-GqD and control mice under HFD (n=9-11). (G) PTT in HFD-fed LysM-GqD and control mice with DCZ (30 µg/kg) stimulation (n=9). (H) Oral glucose tolerance test (O-GTT) in HFD-fed LysM-GqD and control mice with DCZ (30 µg/kg) stimulation (n=6-9). Data represented as mean ± s.e.m. using One-tailed unpaired t-test (A-H); *p < 0.05, **p < 0.01, ***p < 0.001; NS = not significant.

**Supp Figure 3.**
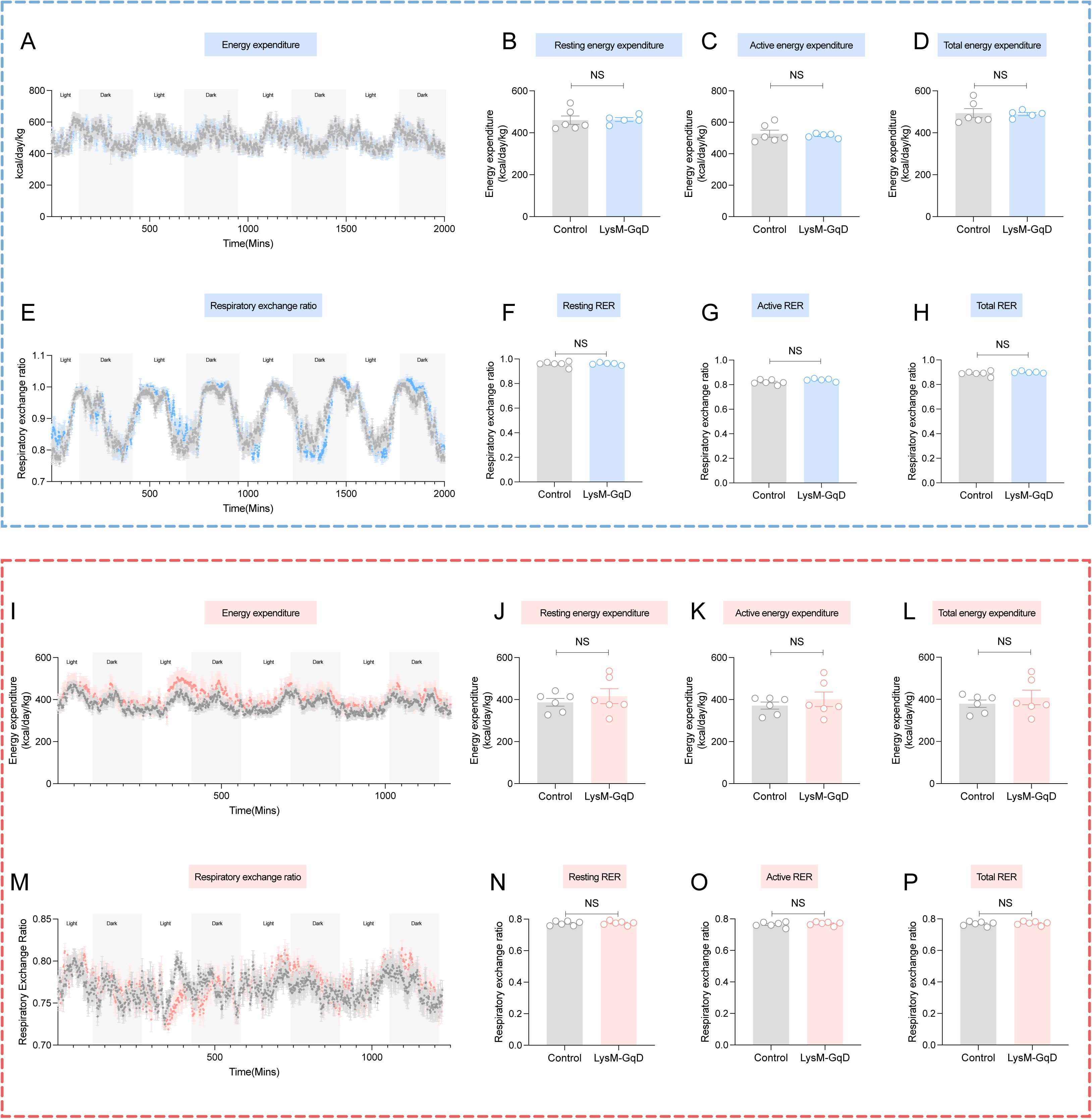
Clodronate liposome treatment effectively depletes islet macrophages. (A) Quantification of F4/80⁺ mRNA expression in islets after clodronate liposome treatment compared with control liposomes (DAPI, blue; F4/80, green), scale-20µM (n = 3). (B) Immunoblot for HA-tag confirming expression of LysM-GqD in macrophages; β-actin serves as loading control. (C) Representative immunofluorescence images of pancreatic islets stained with DAPI (blue) and F4/80 (green) in control versus clodronate liposomes–treated mice. White arrowheads indicate F4/80⁺ macrophages, which are markedly reduced after clodronate treatment (n=3). Data represented as mean ± s.e.m.; ***p < 0.001.

**Supp Figure 4.**
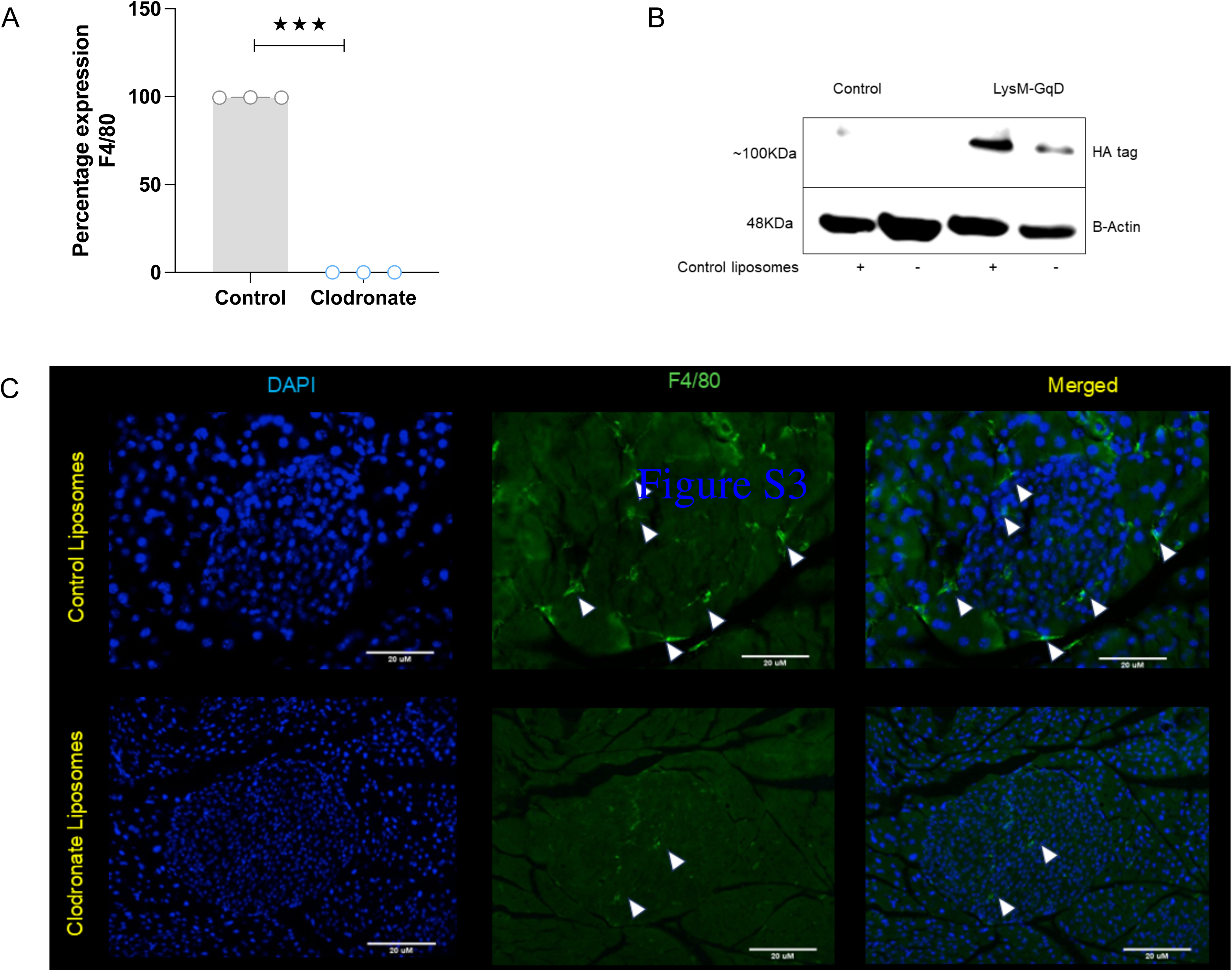
Myeloid Gq Activation Does Not Alter Whole-Body Energy Expenditure or Substrate Utilization both in RC and HFD condition. (A–D) Energy expenditure in RC-fed mice: (A) Time-course of total energy expenditure across light and dark cycles (n=5-6); (B–D) Quantification of resting, active, and total energy expenditure showing no significant differences between genotypes (n=5-6). (E–H) RER in RC-fed mice: (E) RER traces across light and dark cycles; (F–H) Quantification of resting, active, and total RER, indicating comparable substrate utilization between groups. (I–L) Energy expenditure in HFD-fed mice: (I) Time-course of total energy expenditure; (J–L) Quantification of resting, active, and total energy expenditure showing no genotype-dependent differences. (M–P) RER in HFD-fed mice (n=6): (M) RER time-course across light and dark cycles; (N–P) Quantification of resting, active, and total RER, with no significant differences between groups. Data are presented as mean ± s.e.m. using One-tailed unpaired t-test; NS = not significant.

**Supp Figure 5.**
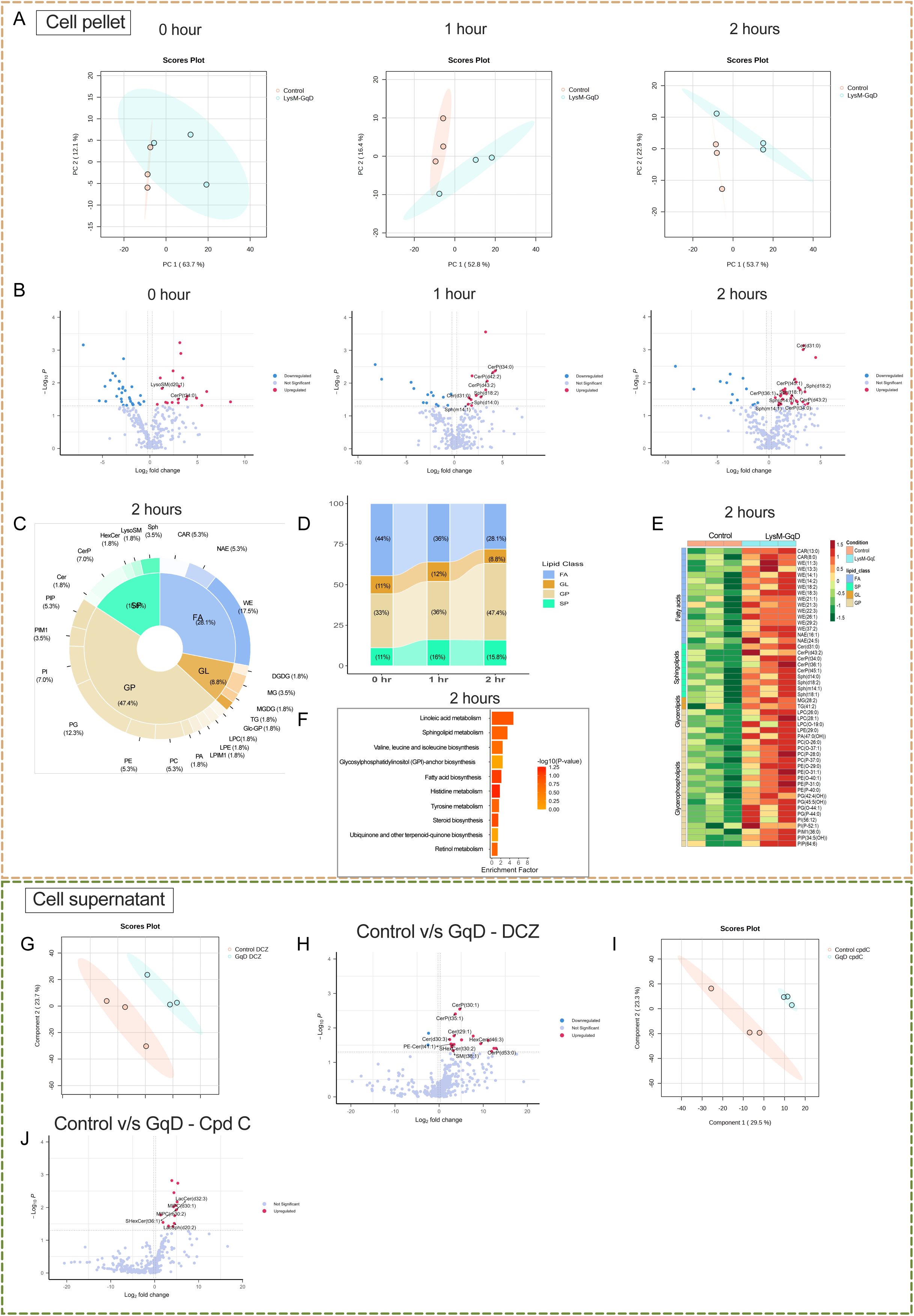
Lipidomic profiling reveals dynamic lipid remodeling in LysM-GqD macrophages. (A–B) Principal component analysis (PCA) (A) and volcano plots (B) of cell pellet lipids at 0, 1, and 2 hours after DCZ stimulation, showing distinct clustering and significant lipid changes at 2 hours. (C) Pie chart illustrating major lipid pathways significantly altered after 2 hours of DCZ treatment. (D) Distribution of lipid classes contributing to differential regulation. (E) Heatmap depicting differentially regulated lipid species at 2 hours in control versus LysM-GqD macrophages. (F) Pathway enrichment analysis identifying lipid metabolic pathways significantly modulated upon Gq activation. (G) PLSDA and (H) volcano plots of lipid changes in cell supernatants comparing control vs LysM-GqD cells following DCZ stimulation. (I) PLSDA and (J) Volcano plot comparing lipid changes between control and LysM-GqD macrophages after Cpd C treatment. All the experiments had 3 biological independent biological replicates and 2 technical replicates for each sample.

**Supp Figure 6.**
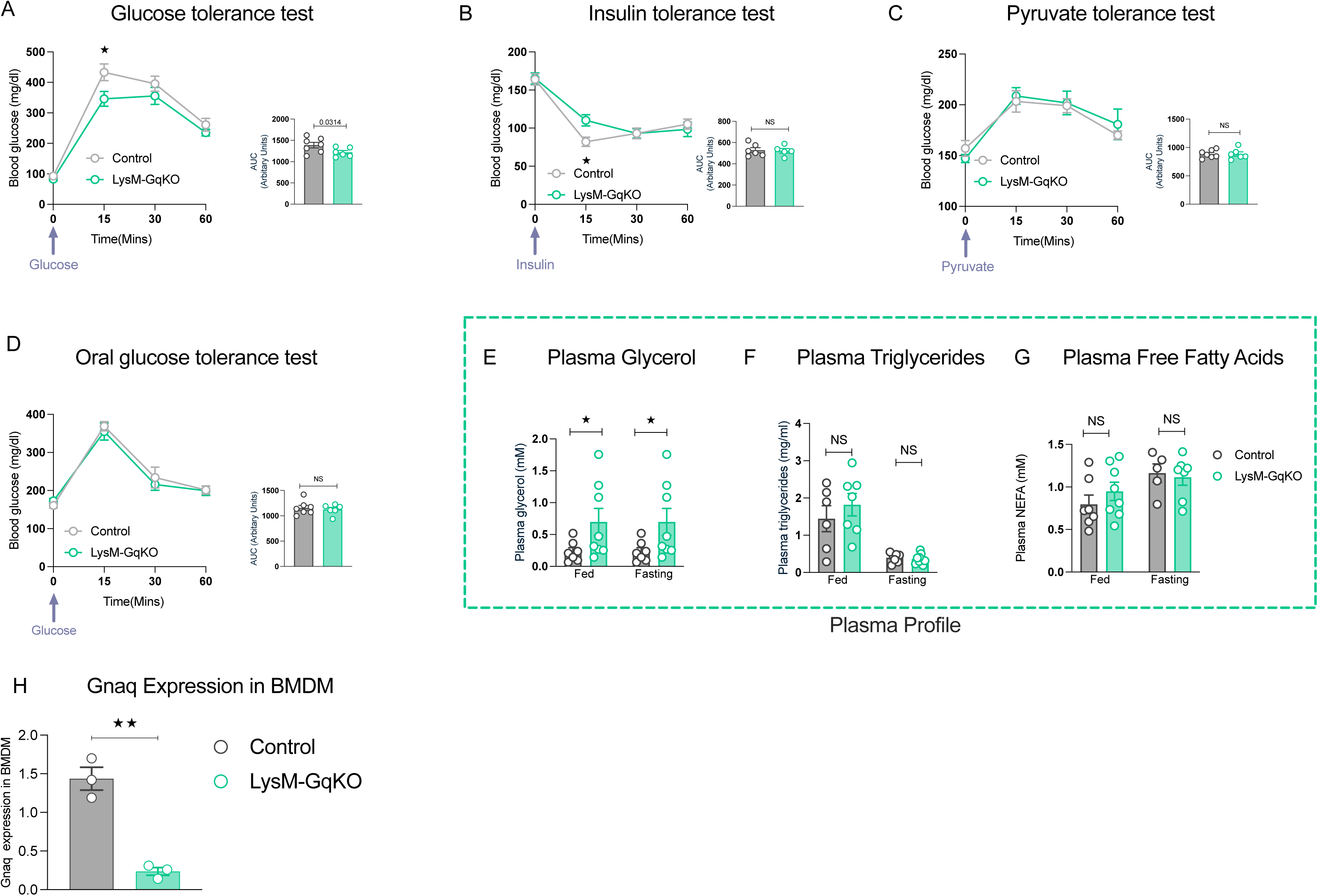
Glucose and lipid metabolic profiling in LysM-Gq KO mice. (A–D) Glucose tolerance test (GTT) (n=6-7) (A), insulin tolerance test (ITT) (n=6-7) (B), pyruvate tolerance test (PTT) (n=6-7) (C), and oral glucose tolerance test (O-GTT) (n=6-7) (D) in control and LysM-Gq KO mice. Knockout mice show modest differences in glucose handling, with largely comparable overall glucose excursions (AUC). (E–G) Plasma profiling in fed and fasted states. Plasma glycerol levels were significantly altered in KO mice (n=7-8) (E), whereas plasma triglycerides (n=7-9) (F) and free fatty acids (n=7-9) (G) remained unchanged. (H) *Gnaq* expression levels in BMDMs from control and LysM-Gq KO mice (n=3). Data represented as mean ± s.e.m. using One-tailed unpaired t-test (A-G); *p < 0.05, **p < 0.01, ***p < 0.001; NS = not significant.

**Supp Figure 7.**
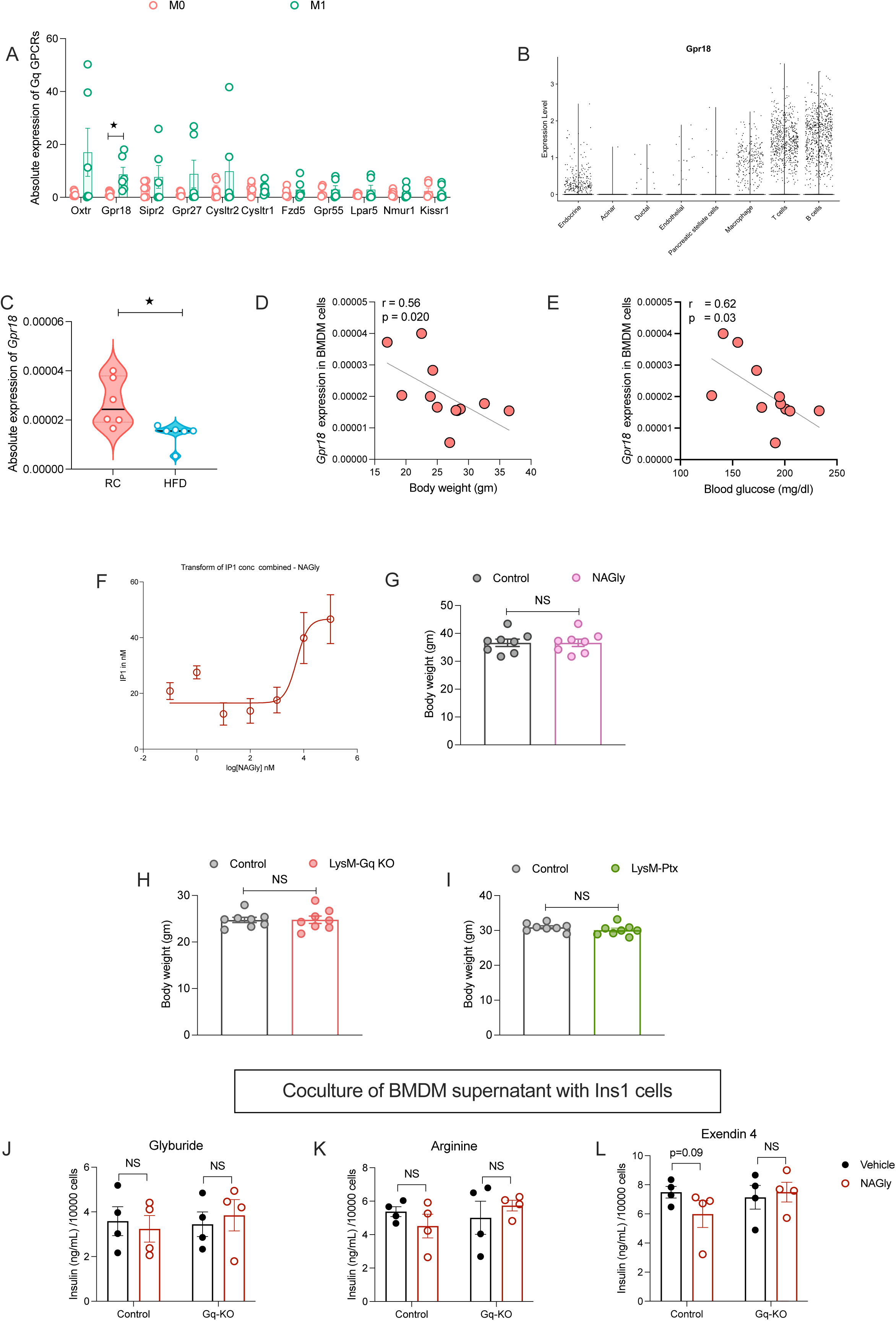
Gpr18 expression and functional assessment in islet macrophages and LysM-Gq KO models. (A) Relative GPCRs expression profiling in murine islet macrophages reveals Gpr18 enrichment. (B) Analysis of publicly available single-cell RNA seq datasets confirms immune cell specific expression of Gpr18 in murine islets. (C–E) Gpr18 expression in macrophages is reduced under HFD conditions (C) and inversely correlates with body weight (D) and blood glucose levels (E). (F) IP-One assay demonstrating NAGly (GPR18 agonist)–induced, dose-dependent IP1 accumulation in DCZ-stimulated GqD macrophages. (G–I) Body weight analysis showing no significant differences between control and NAGly-treated mice (n=8) (G), LysM-Gq KO and control mice (n=8) (H), and LysM-PTX and control mice (n=8) (I). (J–L) Coculture of BMDM and INS-1 cell for insulin secretion assays in control and LysM-Gq KO islets following stimulation with glyburide (n=4) (J), arginine (n=4) (K), or Exendin-4 (n=4) (L). Data represented as mean ± s.e.m. using One-tailed unpaired t-test (A-L); *p < 0.05, **p < 0.01, ***p < 0.001; NS = not significant.

**Supp Figure 8.**
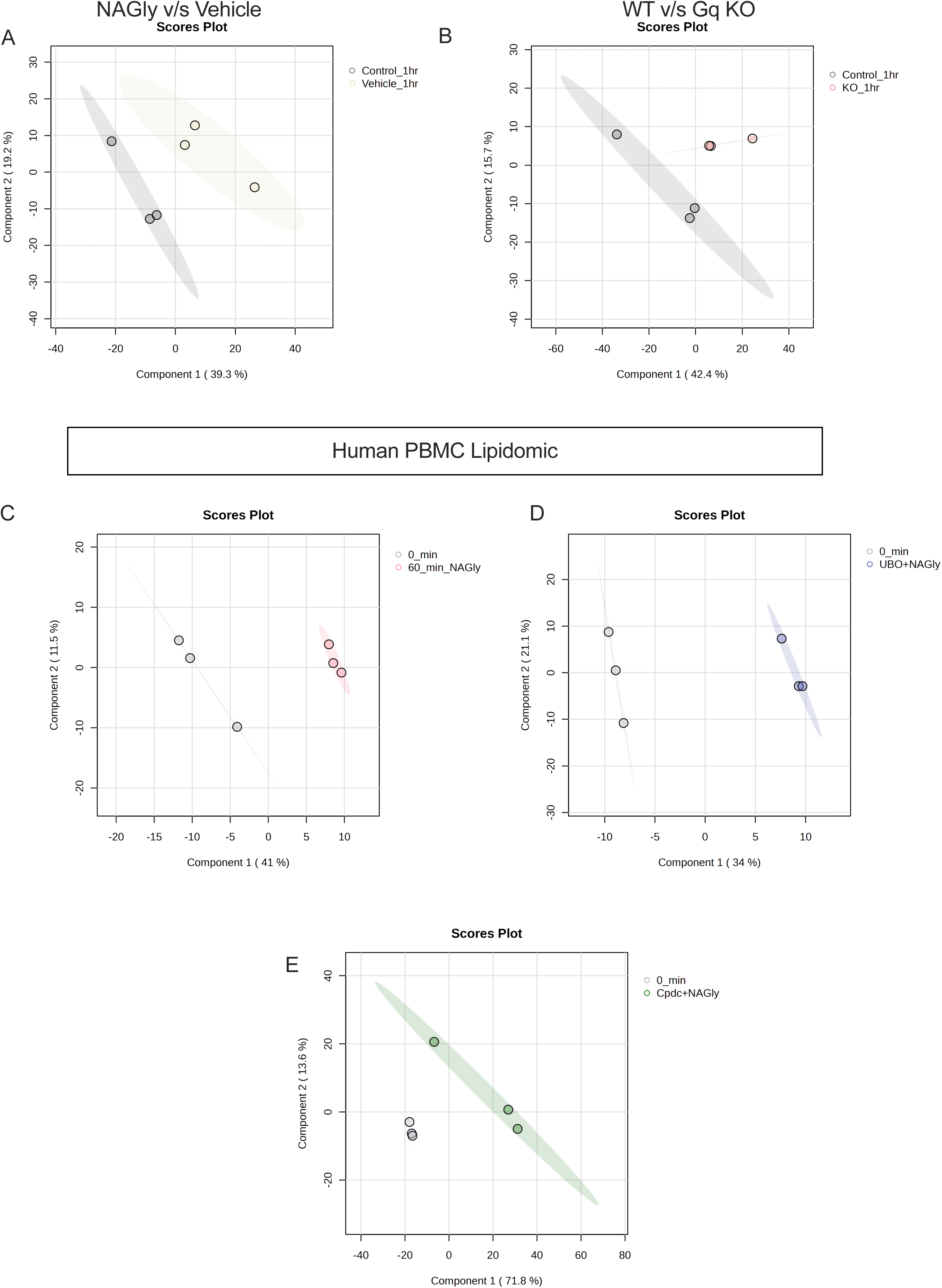
Lipidomic profiling of Control and Gq KO islets following vehicle and NAGly treatments. PLSDA of (A) Vehicle v/s NAGly for 1 hour in WT mice (B) WT v/s NAGly for 1 hour. PL-SDA of (C) 0 hour vs 1 hour NAGly; (D) 0 hour vs UBO+NAGly; (E) 0 hour vs CpdC+NAGly in PBMC.

**Resource Table: 1.**
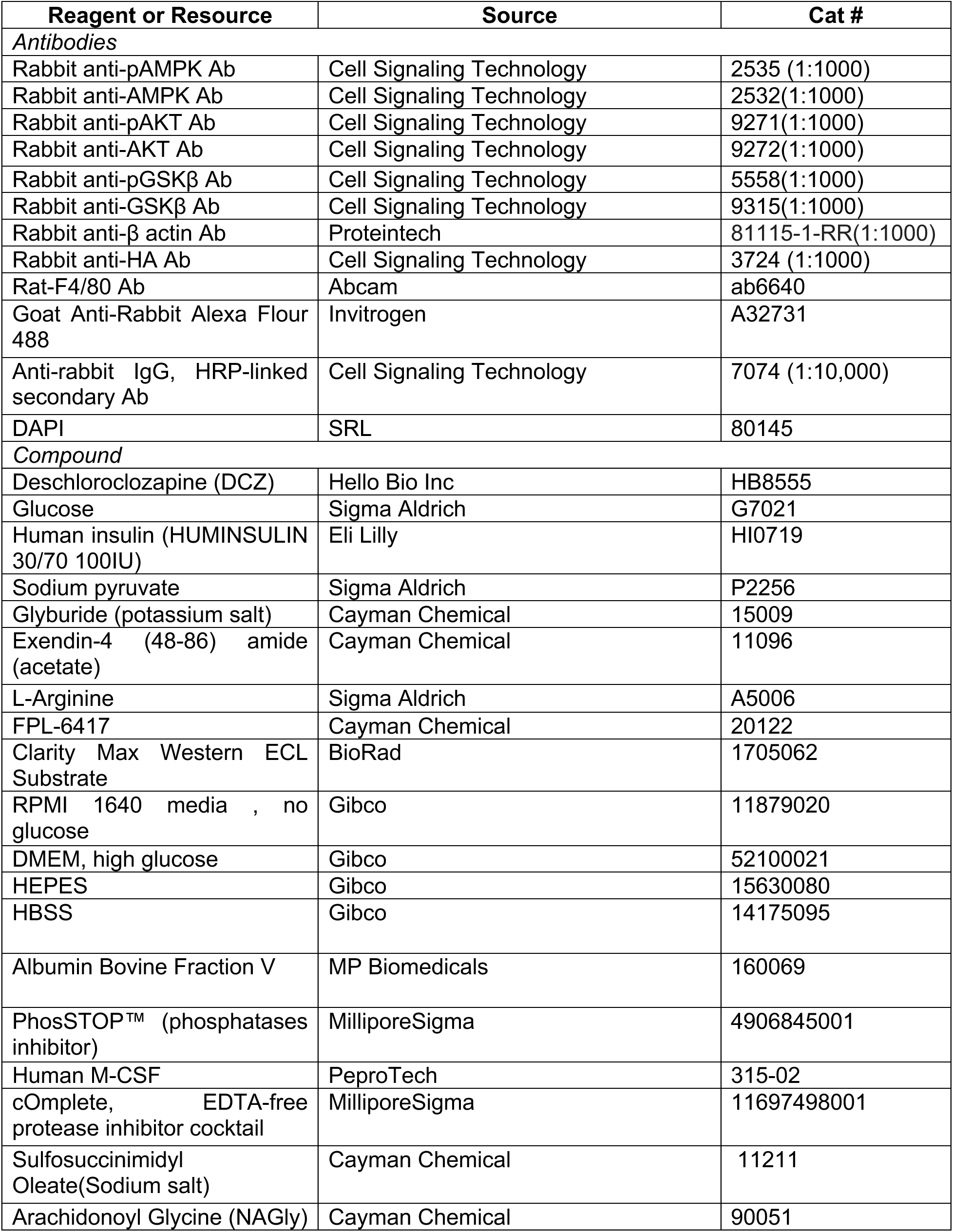
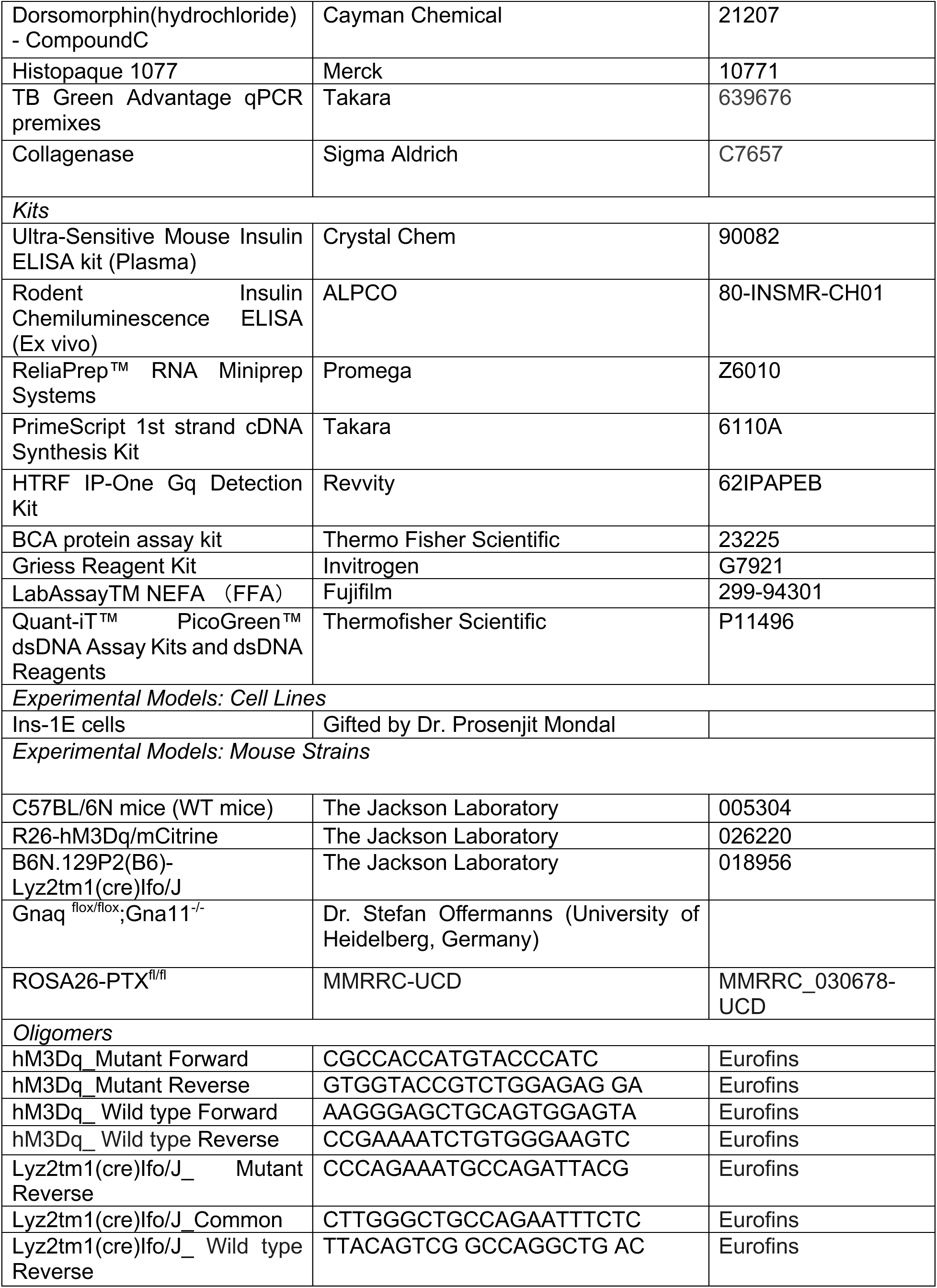
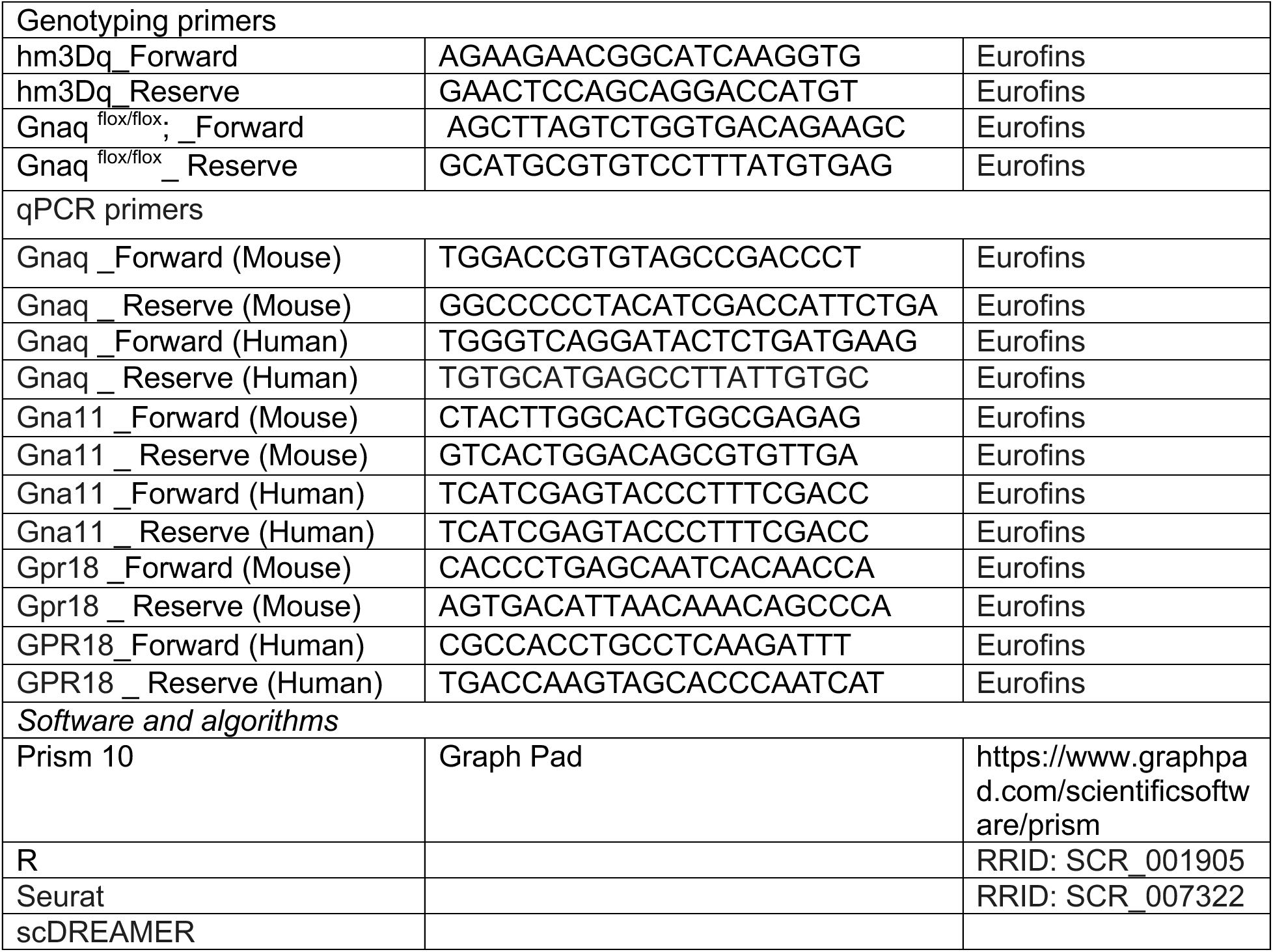

